# Lipid-Driven NRF2 Preconditioning Links Estrogen Deprivation to Hepatic Resilience Against Acetaminophen Injury

**DOI:** 10.64898/2026.11.10.765604

**Authors:** José García-Consuegra, Naroa Goikoetxea-Usandizaga, Ariadna Qin Gil Martín, Ana Sagrera Aparisi, José María Lanza-Arnaiz, Irene González-Recio, Carolina Conter, Raul J. Andrade, M Isabel Lucena, Paolo Grumati, Bruno Ramos Molina, Estela Area, James W Dear, Chris Humphries, Laura Formentini, María Luz Martínez-Chantar, Sara Cogliati

## Abstract

**Background & Aims:** Menopause is associated with a sustained decline in estrogen signaling that profoundly alters hepatic lipid metabolism; however, its impact on the liver response to xenobiotic stress remains poorly understood. Given that acetaminophen (APAP) overdose is a leading cause of drug-induced acute liver failure (DILI) and that sex and age influence susceptibility to hepatotoxicity, we investigated how long-term estrogen deprivation affects the development of APAP-induced liver injury (AP-ILI).

**Methods:** Clinical data from a multicenter cohort of patients admitted following APAP overdose were analyzed to assess susceptibility to AP-ILI. To model the postmenopausal state, 12-week-old female C57BL/6JOlaHsd mice underwent bilateral ovariectomy (OVX) or sham surgery and, six months later, were challenged with acetaminophen (360 or 600 mg/kg). Mechanistic studies included NRF2 silencing, estradiol replacement, and dietary lipid modulation using a high-fat diet. Comprehensive lipidomic profiling was performed to characterize hepatic lipid remodeling, while human liver biopsies were analyzed to validate the expression of key molecular targets.

**Results:** Women ≥50 years of age showed a lower risk of severe AP-ILI, a phenotype recapitulated in OVX mice. Long-term estrogen deprivation induced hepatic lipid remodeling characterized by larger lipid droplets and enrichment in saturated neutral lipids, leading to sustained NRF2 activation. Enhancement of lipid remodeling with high-fat diet increased NRF2 signaling and hepatoprotection, whereas estradiol replacement reversed these effects. Consistent with these findings, women over 50 years of age exhibited increased hepatic expression of *NRF2* and its target genes.

**Conclusions:** Long-term estrogen deprivation confers protection against AP-ILI through extensive hepatic lipid remodeling, characterized by lipid droplet expansion and altered neutral lipid composition, which promotes sustained NRF2 activation.

**IMPACT AND IMPLICATIONS:** These findings identify menopause-associated metabolic remodeling as a previously unrecognized determinant of resistant to drug-induced liver injury. By linking long-term estrogen deprivation to sustained NRF2 activation through hepatic lipid remodeling, this study uncovers a novel endogenous hepatoprotective mechanism that may contribute to the lower risk of severe AP-ILI observed in postmenopausal women. Beyond acetaminophen toxicity, these results suggest that age-and sex-dependent changes in hepatic lipid metabolism could broadly influence liver resilience to xenobiotic and oxidative stress, opening new opportunities for risk stratification and NRF2-targeted therapeutic strategies in DILI.

**HIGHLIGHTS:**

- Women ≥50 years exhibited a lower risk of AP-ILI, a phenotype recapitulated in OVX mice.
- OVX induced hepatic lipid remodeling, with increased saturated lipid species and enlarged lipid droplets.
- Hepatic lipid remodeling promoted sustained activation of the NRF2 pathway.
- Estradiol supplementation reversed NRF2 activation and the hepatoprotective phenotype in OVX mice.
- Women ≥50 years have increased hepatic expression of NRF2 supporting the translational relevance of these findings.

**Graphical abstract:** 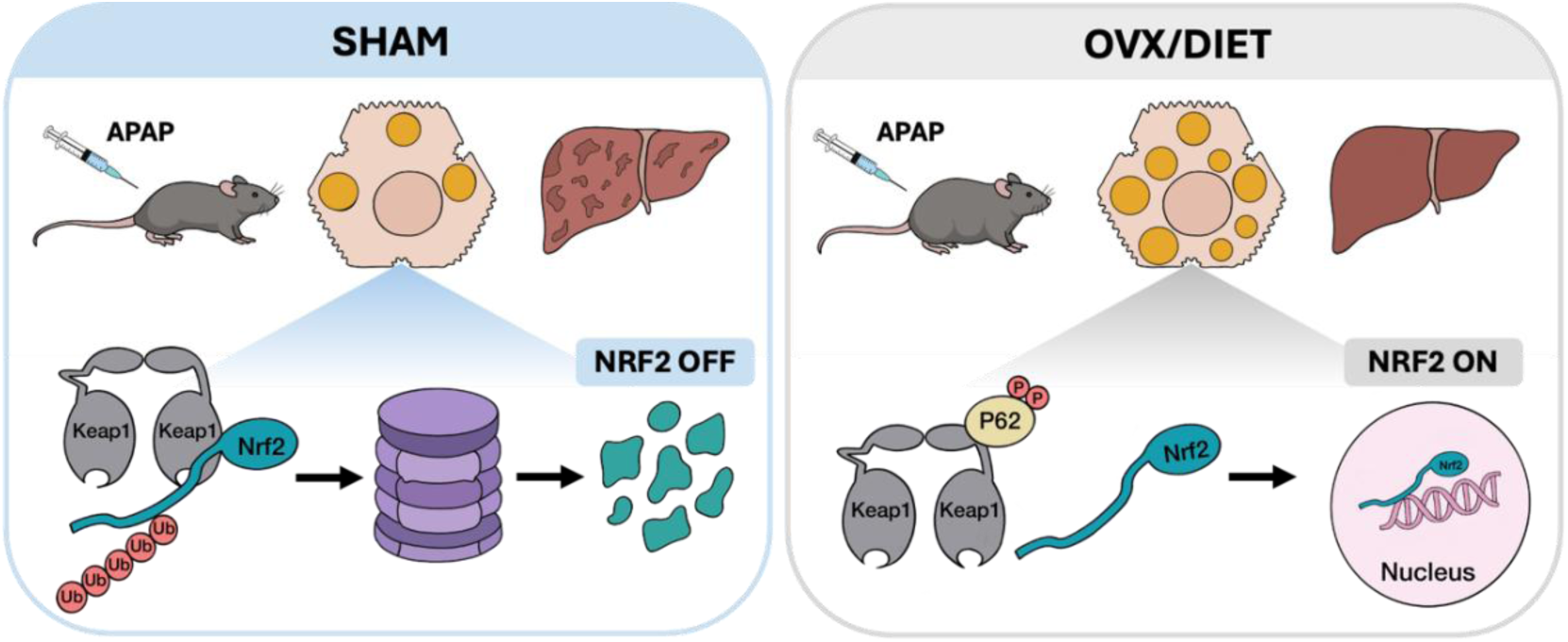

## INTRODUCTION

Menopause is characterized by the permanent cessation of ovarian function and a marked decline in estrogen levels. This endocrine transition is accompanied by profound metabolic alterations that increase susceptibility to metabolic dysfunction-associated steatotic liver disease (MASLD) [1,2] and a broad spectrum of cardiometabolic disorders [3–7]. These changes include alterations in lipid metabolism, cholesterol homeostasis, and redox balance, all of which contribute to metabolic dysfunction.

In the liver, estrogen signaling is a major regulator of lipid homeostasis and coordinates the expression of genes involved in lipid synthesis, storage, and utilization throughout the estrous cycle [8]. Consistent with this role, disruption of estrogen signaling promotes lipogenic pathways while suppressing lipid catabolism, thereby favoring lipid droplet accumulation, hepatic inflammation, and fibrosis, and contributing to the dyslipidemic profile commonly observed after menopause [9–11]. However, despite extensive knowledge regarding the metabolic consequences of estrogen deficiency, considerably less is known about how chronic estrogen deprivation affects the hepatic response to acute toxic stress.

Drug-induced liver injury (DILI) remains a major clinical challenge and is one of the leading causes of serious liver injury and acute liver failure worldwide [12,13]. Among the different forms of DILI, acetaminophen (APAP) overdose represents the prototypical model of intrinsic hepatotoxicity and continues to be a leading cause of drug-induced acute liver failure in Western countries [14,15].

Sex and age are increasingly recognized as important modifiers of DILI susceptibility and outcome, although their effects appear to be drug-specific and remain incompletely understood. Women are overrepresented among patients with APAP-induced acute liver injury and acute liver failure, and female sex has been associated with greater clinical severity in large prospective cohorts [16]. More broadly, sex-related differences in DILI vary with age, raising the possibility that ovarian hormone status influences the hepatic response to xenobiotic injury [17]. Nevertheless, whether long-term estrogen deprivation directly modifies susceptibility to APAP-induced liver injury (AP-ILI), and through which mechanisms, remains unknown.

Accumulating evidence indicates that lipid droplet formation is not invariably detrimental [18]. Beyond its well-established role in metabolic disease, lipid remodeling is increasingly recognized as a dynamic adaptive response that protects cells from environmental, metabolic, and oxidative stress [19]. Such adaptations have also been linked to activation of the NRF2 antioxidant pathway [20–22]. However, it remains unclear whether menopause-associated hepatic lipid remodeling engages these protective mechanisms and contributes to resistance against DILI.

Here, we identify a previously unrecognized lipid-dependent adaptive program triggered by long-term estrogen deprivation that promotes hepatic resilience to APAP-induced liver injury. We show that postmenopausal women exhibit a lower risk of AP-ILI following overdose and that prolonged ovarian hormone deprivation in mice induces the accumulation of saturated neutral lipids, enlargement of lipid droplets, and sustained activation of the NRF2 pathway. Together, these adaptations establish a preconditioned hepatic state that enhances resistance to APAP-induced oxidative stress. Nutritional interventions that further promoted lipid remodeling augmented NRF2 activation and hepatoprotection, whereas estradiol replacement reversed these effects. Moreover, liver samples from women over 50 years of age displayed increased expression of NRF2 and its target genes, supporting the translational relevance of our findings.

Collectively, our data identify chronic estrogen deprivation as a driver of lipid-dependent hepatic preconditioning and reveal sustained NRF2 activation as a previously unrecognized mechanism conferring resilience to APAP-induced toxic liver injury.

## RESULTS

### Postmenopausal women and ovariectomized female mice are protected from APAP-induced liver injury (AP-ILI)

To assess the risk of AP-ILI, we analyzed a multicenter cohort including all patients admitted following APAP overdose across three hospitals (Table 1). Multivariable analysis using Firth-penalized logistic regression revealed that, after adjustment for paracetamol concentration and relevant biochemical and demographic variables, women aged ≥50 years exhibited a significantly lower risk of severe liver injury than women <35 years (adjusted odds ratio [aOR] 0.18; 95% CI 0.04-0.75; p = 0.018). A similar trend was observed in men aged ≥50 years (aOR 0.28; 95% CI 0.07-1.11; p = 0.07), whereas men <35 years showed no significant difference relative to the reference group (aOR 0.75; p = 0.35) (Fig. 1A).

**Table 1.** Demographic and clinical characteristics of the patients included in the study.

| Cohort | Total N | Severe Hepatitis Events, n (%) | Age, years, Median (IQR) | Presentation ALT, U/L, Median (IQR) | Peak ALT, U/L, Median (IQR) | Presentation Paracetamol, mg/L, Median (IQR) | Hospital Mortality, n (%) |
|---|---|---|---|---|---|---|---|
| **Men ≥50 years** | 524 | 15 (2.9%) | 57 (53–63) | 24 (16–42) | 29 (19–55) | 26 (6–82) | 5 (1.0%) |
| **Men <35 years** | 1,163 | 21 (1.8%) | 25 (20–29) | 22 (15–36) | 25 (17–43) | 44 (9–102) | 1 (0.1%) |
| **Women <35 years<br>(Baseline)** | 2,663 | 35 (1.3%) | 21 (18–27) | 15 (12–22) | 17 (13–26.5) | 56 (16–114) | 2 (0.1%) |
| **Women ≥50 years** | 762 | 14 (1.8%) | 56 (52–62) | 21 (14–32.8) | 24 (16–41) | 55.5 (13–122) | 5 (0.7%) |
| **Overall Total** | **5,112** | **85 (1.7%)** | **26 (20–50)** | **18 (13–29)** | **21 (14–35)** | **50 (12–111)** | **13 (0.3%)** |

**Figure 1.**
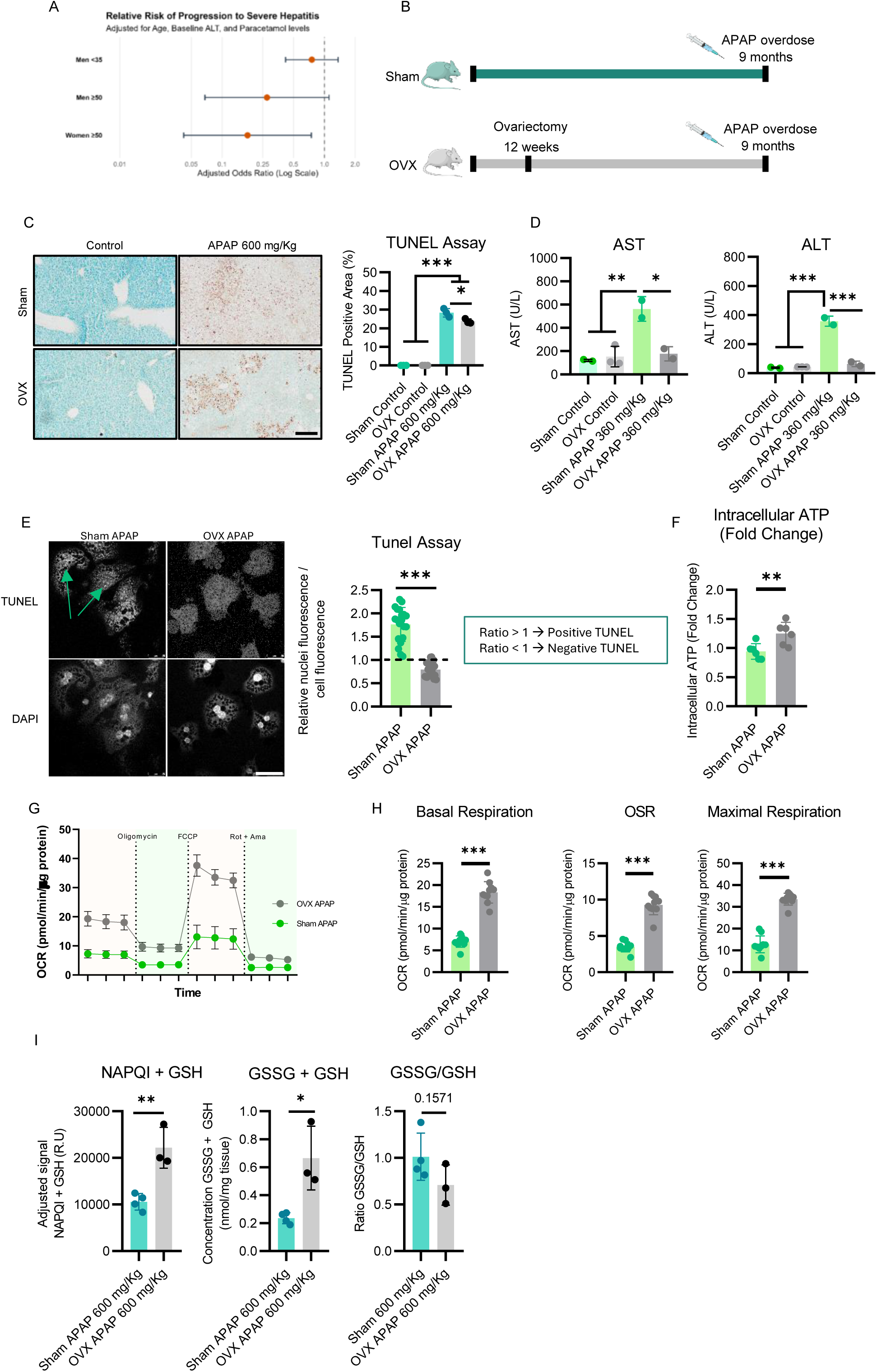
Postmenopausal women and ovariectomized (OVX) female mice are protected against APAP-induced liver injury and preserve mitochondrial function. (A) Multivariable Firth-penalized logistic regression analysis showing the adjusted odds ratio for progression to severe hepatitis after APAP overdose. Fertile women (<35 years old) were used as the reference group (OR = 1). (B) Experimental design and study workflow. (C) Representative TUNEL staining of liver sections and quantification of TUNEL-positive areas (n = 3 animals per group). Brown staining indicates necrotic tissue, whereas green staining indicates viable tissue (scale bar = 500 μm). (D) Serum AST and ALT activities in control and APAP-treated mice (360 mg/kg; n = 2-3 animals per group). (E) Representative TUNEL staining of primary hepatocytes isolated 24 h after in vivo APAP administration (360 mg/kg), together with its quantification (scale bar = 50 μm). TUNEL positivity was determined as the ratio between nuclear TUNEL signal and background cellular signal, with values >1 considered positive. Each dot represents an individual cell (n = 20-21 cells from 3 technical replicates). (F) Intracellular ATP levels in primary hepatocytes isolated from control and APAP-treated mice (n = 6 technical replicates). (G) Representative oxygen consumption rate (OCR) profiles of primary hepatocytes isolated from control and APAP-treated mice. (H) Quantification of basal respiration, oligomycin-sensitive respiration (OSR), and maximal respiration (n = 10 technical replicates). (I) Hepatic NAPQI-GSH adducts, total glutathione (GSH + GSSG) levels, and GSSG/GSH ratio measured in whole liver tissue (n = 3-4 animals per group). Data are presented as mean ± SD. Statistical significance was assessed using an unpaired Student’s t-test in (E), (F), (H), (I) and one-way ANOVA followed by Tukey’s multiple-comparison test in (C) and (D) *p < 0.05, **p < 0.01, ***p < 0.001.

To investigate the mechanisms underlying this age-dependent protection, we employed a mouse model of long-term estrogen deficiency that mimics postmenopause. Female C57BL/6J mice underwent ovariectomy (OVX) or sham surgery and were challenged with APAP (600 mg/kg) six months later (Fig. 1B). OVX mice exhibited improved survival (Fig. S1A) and reduced hepatocellular death compared with sham-operated controls (Fig. 1C). Because this dose induced fulminant liver injury and early mortality, subsequent mechanistic studies were performed using a lower APAP dose (360 mg/kg). Under these conditions, OVX mice displayed markedly lower serum ALT and AST levels and reduced hepatocyte death, as determined by TUNEL staining, confirming protection against AP-ILI (Fig. 1D, E). Because APAP hepatotoxicity is closely linked to mitochondrial dysfunction and ATP depletion, we assessed these parameters in sham and OVX hepatocytes after APAP exposure. OVX hepatocytes maintained higher ATP levels and displayed greater basal, ATP-linked, and maximal respiration than sham cells (Fig. 1F-G). Consistently, APAP induced severe mitochondrial damage in sham livers, including mitochondrial swelling, cristae disruption, and loss of structural integrity, whereas mitochondrial morphology was largely preserved in OVX mice (Fig. S1B-D). Accordingly, OVX livers retained higher levels of the cristae-shaping protein OPA1^L^ (Fig. S1E) which is associated with cristae integrity and mitochondrial respiration [23]. Together, these findings indicate that long-term estrogen deprivation enhances resistance to APAP-induced liver injury and preserves mitochondrial bioenergetics and integrity.

Given the preserved mitochondrial function and reduced liver injury in OVX mice, we next assessed APAP bioactivation and detoxification. APAP overdose causes excessive CYP2E1-dependent NAPQI formation, overwhelming glutathione-mediated detoxification and triggering oxidative stress and mitochondrial dysfunction. After administration of 600 mg/kg APAP, OVX animals have increased levels of NAPQI-GSH and total levels of GSSG+GSH while, the GSSG/GSH ratio was lower, indicating enhanced redox buffering capacity (Fig. 1I). Consistent with this, OVX livers showed increased expression of *Gsr, Gsta*, and *Gclc* (Fig. S1F), involved in glutathione turn-over. Together, these findings indicate that the reduced oxidative stress observed in OVX mice following APAP overdose could be driven at least partially by enhanced glutathione-dependent detoxification.

### Long-term estrogen deficiency reshapes hepatic lipid composition and preserves membrane lipid homeostasis during APAP injury

We hypothesized that the lipid remodeling induced by chronic estrogen deprivation contributes to the protection against AP-ILI observed in OVX mice. Consistent with a shift toward lipid storage, OVX livers displayed reduced expression of β-oxidation genes and increased expression of lipogenic genes (Fig. 2A), accompanied by enlarged lipid droplets despite unchanged hepatic TAG content and absence of overt steatosis (Fig. 2B-D). Lipidomic analysis revealed a marked reduction in saturated cholesteryl esters, selective enrichment of saturated TAG species, and remodeling of arachidonic acid-containing phospholipids (Fig. 2E-G), indicating profound lipid reorganization rather than simple lipid accumulation.

**Figure 2.**
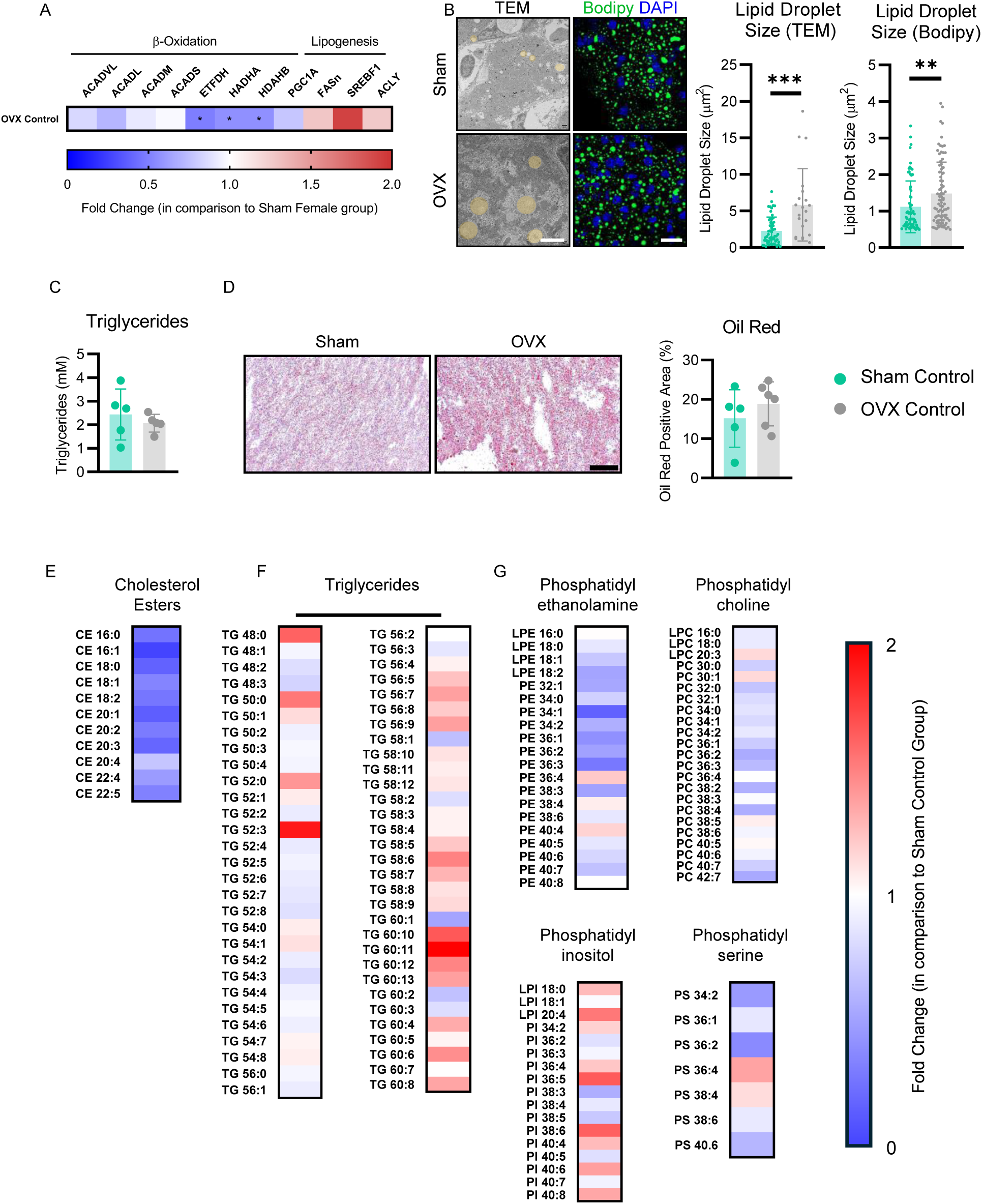
Ovariectomy induces hepatic lipid remodeling and promotes lipid droplet expansion. (A) Relative hepatic mRNA expression of the indicated genes, expressed as fold change relative to the Sham female group (n = 3-4 animals per group). (B) Representative transmission electron microscopy (TEM) images of liver tissue. Lipid droplets are highlighted in yellow (scale bar = 5 μm). Representative BODIPY staining of neutral lipids (green) in liver sections with DAPI nuclear counterstaining (blue) (scale bar = 20 μm). For TEM analyses, 7-20 randomly selected lipid droplets from 3 independent animals were quantified. For BODIPY analyses, 14 randomly selected lipid droplets from 5-6 animals per group were quantified. For both quantifications, a similar number of lipid droplets was quantified for each animal. (C) Hepatic triglyceride content measured in whole liver tissue (n = 5 animals per group). (D) Representative Oil Red O staining of liver sections and corresponding quantification in Sham and OVX mice (n = 5-7 animals per group). Scale bar = 500 μm. (E-G) Heatmaps showing the relative abundance of the indicated lipid species, expressed as fold change relative to the Sham female group. Data in panels (B-D) are presented as mean ± SD. Statistical significance was assessed using unpaired Student’s t-tests. **p < 0.001.

We next examined how this lipid phenotype influenced the response to APAP. In fertile females, APAP triggered extensive lipid remodeling characterized by sphingolipid alterations, accumulation of PUFA-containing neutral lipids, and depletion of membrane phospholipids (Fig. 3A-H). While OVX mice displayed a similar redistribution of lipids toward neutral lipid pools, membrane phospholipid loss was markedly attenuated and accompanied by increased levels of antioxidant lipid species, including CoQ9 and PEos (Fig. 3A-I). Notably, enlarged lipid droplets were maintained after APAP challenge (Fig. 3J). Together, these findings identify a distinct lipid remodeling program in OVX livers characterized by enlarged lipid droplets, preservation of membrane phospholipids, and enrichment of antioxidant lipids, which may contribute to enhanced resistance to AP-ILI.

**Figure 3.**
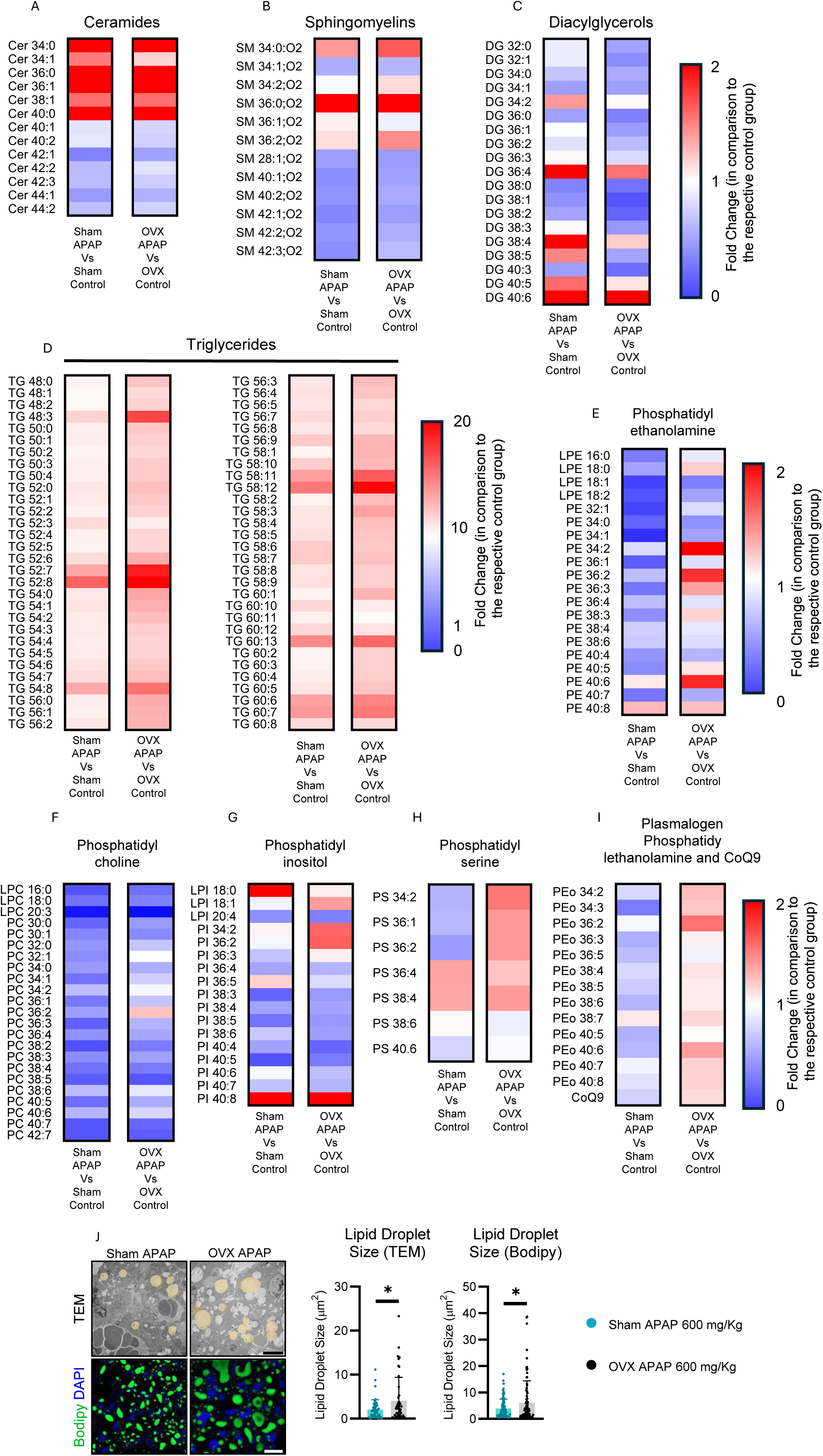
APAP exposure remodels the hepatic lipidome and modifies lipid droplet morphology in OVX mice. (A-I) Heatmaps showing the relative abundance of the indicated lipid species, expressed as fold change relative to the indicated experimental group. (J) Representative transmission electron microscopy (TEM) and BODIPY images of liver tissue and corresponding quantification of lipid droplet size. For TEM, scale bar = 5 μm and a total of 15 randomly selected lipid droplets from 3 independent animals were quantified. For BODIPY staining scale bar = 20 μm, 15 randomly selected lipid droplets from 6 animals per group were quantified. Data are presented as mean ± SD. Statistical significance was assessed using unpaired Student’s t-tests. **p < 0.01, ***p < 0.001.

### Nutritional lipid remodeling promotes hepatic resistance to APAP-induced injury

We next examined whether nutritional modulation of hepatic lipid metabolism could recapitulate the resistance to AP-ILI observed in OVX females.

As an initial approach, primary hepatocytes isolated from fertile sham females were cultured in methionine-and choline-deficient (MCD) medium, a well-established model of hepatocellular lipid stress [24]. Under these conditions, cells displayed significantly reduced APAP-induced cell death compared with control cultures (Fig. S2A, B), suggesting that lipid remodeling per se may confer resistance to APAP toxicity.

To determine whether this phenomenon could also be reproduced in vivo, fertile female mice were fed a high-fat diet (HFD; 60% fat for 5 consecutive days), a dietary intervention previously shown to increase hepatic lipid content and promote lipid droplet enlargement [25,26] (Fig. 4A). Consistent with the in vitro findings, HFD-fed mice exhibited reduced hepatocellular death following APAP overdose, as assessed by TUNEL staining (Fig. 4B), together with improved survival compared with animals maintained on standard chow (Fig. S1A). Notably, HFD-fed mice also displayed enlarged hepatic lipid droplets after APAP administration (Fig. 4C), closely resembling the phenotype observed in OVX mice. Together, these findings support a direct association between hepatic lipid droplet remodeling and protection against AP-ILI.

**Figure 4.**
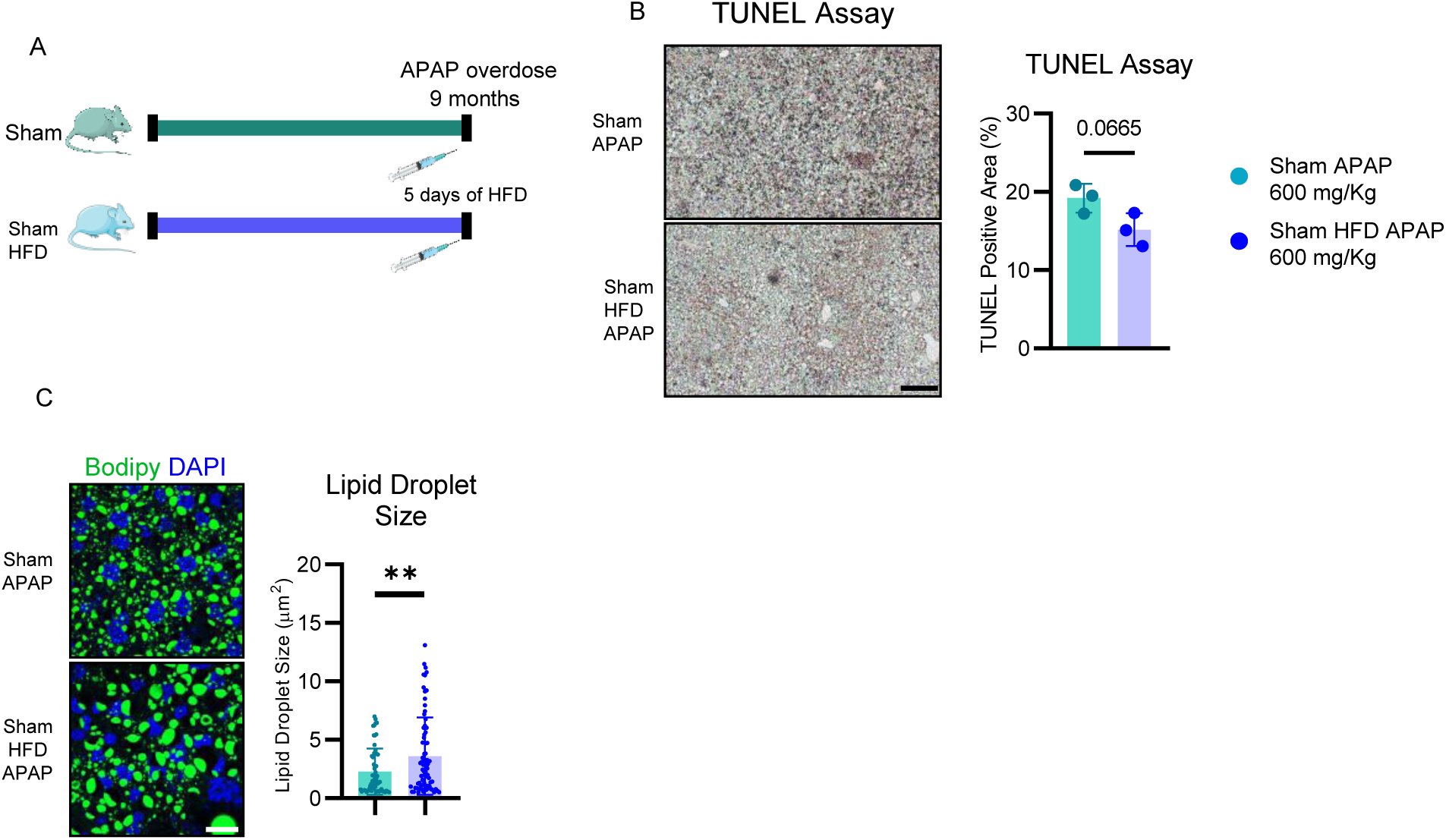
Dietary interventions modulate susceptibility to APAP-induced liver injury in fertile and OVX female mice. High-fat diet enhances resistance to APAP in fertile females, whereas calorie restriction reduces protection in OVX females. (A) Schematic representation of the experimental design and study workflow. (B) Representative TUNEL staining of liver sections and quantification of TUNEL-positive necrotic areas (n = 3 animals per group). Brown staining indicates necrotic tissue, whereas green staining indicates viable tissue. (C) Representative BODIPY staining of neutral lipids in liver sections and corresponding quantification of lipid droplet size. 15 randomly selected lipid droplets per animal were quantified from 4-5 animals per group. Scale bar in TUNEL staining corresponds to 500 μm, while in BODIPY stained liver sections, scale bar corresponds to 20 μm. Data are presented as mean ± SD. Statistical significance was assessed using unpaired Student’s t-tests. *p < 0.05, **p < 0.01, ***p < 0.001.

### Long-term estrogen deficiency activates the hepatic p62–KEAP1–NRF2 antioxidant pathway

To identify potential molecular determinants of this phenotype, we performed unbiased proteomic profiling of APAP-treated sham and OVX livers. This analysis identified increased expression of the NRF2 target proteins HO-1, p62/SQSTM1, and SRXN1 in OVX mice (Fig. 5A). These findings were confirmed by RT-qPCR and western blot analyses (Fig. 5B, C), supporting enhanced NRF2 signaling as a key mechanism underlying the protective phenotype.

**Figure 5.**
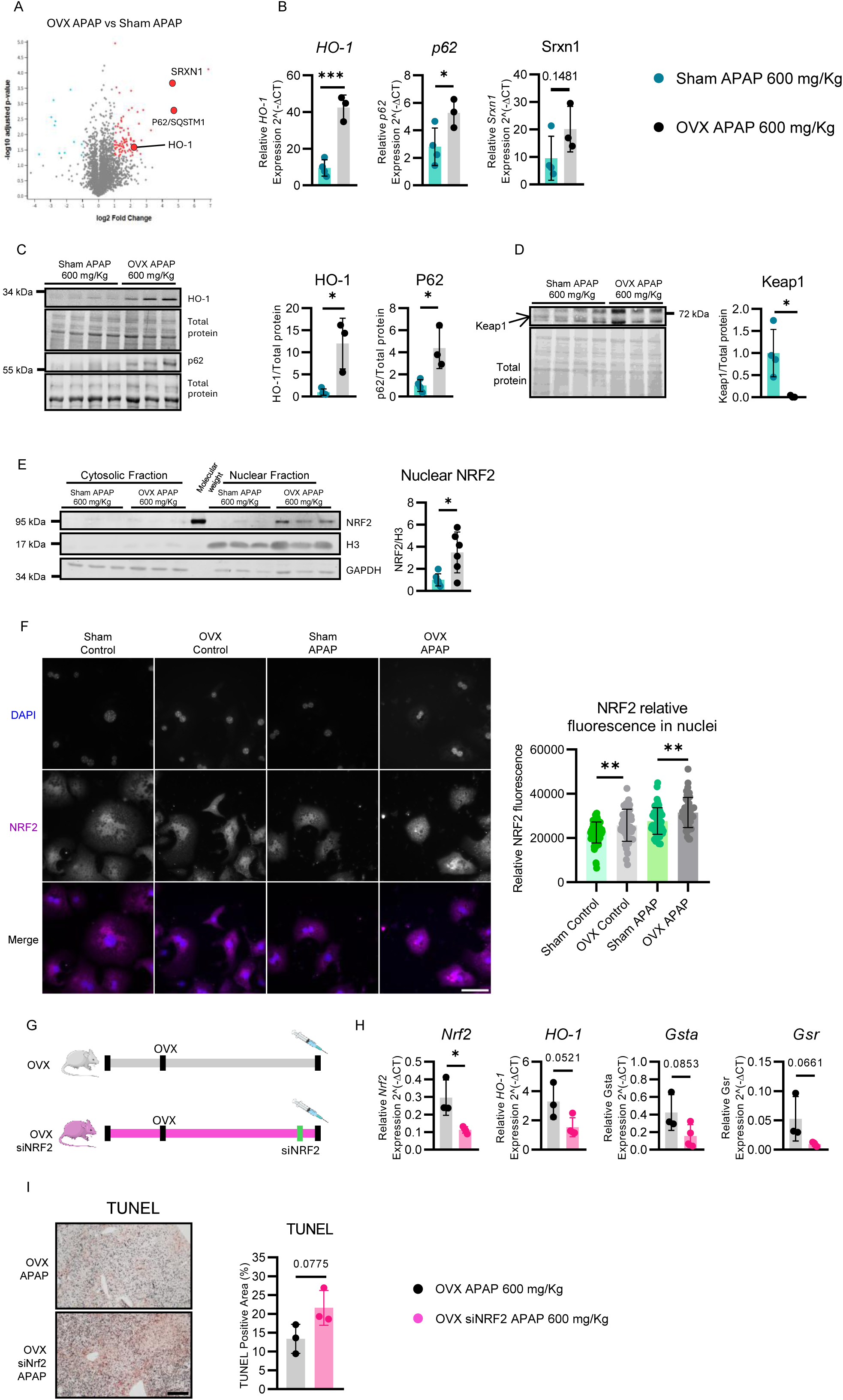
Ovariectomy promotes antioxidant responses and NRF2 stabilization following APAP overdose. (A) Volcano plot of the proteomic analysis comparing OVX and Sham mice following APAP overdose. Upregulated proteins are shown in red and downregulated proteins in blue. (B) Relative hepatic mRNA expression of Hmox1 (HO-1), Sqstm1 (p62), and Srxn1, normalized to Arp expression (n = 3-4 animals per group). (C) Representative immunoblots of HO-1, p62 in whole liver lysates and the relative densitometric quantification normalized to total protein content, with each dot representing an individual animal (n = 3-4 animals per group). (D) Representative immunoblot of KEAP1 in whole liver lysates and the relative densitometric quantification normalized to total protein content, with each dot representing an individual animal (n = 3-4 animals per group). (E) Representative immunoblots showing NRF2 subcellular localization together with densitometric quantification of nuclear NRF2 levels normalized for to the H3 protein level (n = 6 animals per group). (F) Representative immunofluorescence images of NRF2 in primary hepatocytes isolated from control and APAP-treated mice, together with quantification of nuclear NRF2 fluorescence intensity (n = 43-77 cells from 3 different technical replicates) (scale bar = 50 μm). (G) Schematic representation of the experimental design and study workflow. (H) Relative hepatic mRNA expression of *Nrf2*, *Hmox1* (HO-1), *Gsta*, and *Gsr*, normalized to Arp expression (n = 3 animals per group). (I) Representative TUNEL staining of liver sections and quantification of TUNEL-positive necrotic areas (n = 3 animals per group). Brown staining indicates necrotic tissue (scale bar = 500 μm). Each dot represents an individual animal. Data are presented as mean ± SD. Statistical significance was assessed using unpaired Student’s t-tests in (B), (C), (D), (E), (F), (H), (I). For panel (F), comparisons were performed between Sham control versus OVX control and Sham APAP versus OVX APAP groups. *p < 0.05, **p < 0.01, ***p < 0.001.

Consistently OVX livers displayed reduced KEAP1 following APAP administration (Fig. 5D). Subcellular fractionation of liver tissue (Fig. 5E) and immunofluorescence analysis of primary hepatocytes (Fig. 5F) further demonstrated increased nuclear accumulation of NRF2 in OVX mice compared with sham controls under both basal and APAP-treated conditions.

To determine whether NRF2 mediates the protective phenotype of OVX females, we performed siRNA-mediated knockdown experiments. *Nrf2* silencing in primary hepatocytes significantly increased APAP-induced cell death in both sham and OVX groups (Fig. S2A-C). In vivo, we administered *Nrf2*-targeting or control siRNA to OVX mice 24 h before APAP challenge (Fig. 5G). *Nrf2* expression was reduced and its target genes (*HO-1*, *Gsta*, *Gsr*, and *Gclc*) showed a consistent downward trend following siRNA treatment, indicating partial suppression of hepatic NRF2 signaling, although residual activity remained (Fig. 5H). Following APAP overdose, *Nrf2*-silenced OVX mice exhibited increased hepatocellular death compared with control animals (Fig. 5I). While these differences did not reach statistical significance (p > 0.05), reflecting the incomplete silencing achieved by systemic siRNA delivery, the direction and magnitude of the response were remarkably consistent with the results obtained in primary hepatocytes (Fig. S2B), confirming NRF2 as effector of AP-ILI protection.

Given the extensive lipid remodeling induced by long-term estrogen deprivation, we hypothesized that lipid-derived stress signals engage NRF2 and promote a preconditioned antioxidant state. Consistent with this hypothesis, nutritional induction of lipid remodeling through short-term HFD feeding, which recapitulated the AP-ILI-resistant phenotype observed in OVX mice, also increased the expression of the NRF2 target proteins HO-1 and p62 (Fig. S3A). Together, these findings suggest that lipid remodeling acts as an upstream adaptive response that sustains NRF2 activation, thereby establishing a preconditioned hepatic state that enhances resistance to AP-ILI.

### Estradiol replacement reverses the hepatoprotective phenotype of long-term estrogen deficiency

While nutritional interventions identify that metabolic remodeling can modulate NRF2 activity and susceptibility to AP-ILI, the upstream hormonal signal responsible for establishing this adaptive phenotype in OVX females remained to be defined. Because ovariectomy markedly reduces circulating ovarian hormones, particularly 17β-estradiol (E2), we next asked whether restoration of estradiol signaling was sufficient to reverse the hepatoprotective state induced by long-term estrogen deficiency. OVX females received E2 supplementation (1.25 µg/day) for 15 consecutive days starting two months after ovariectomy, prior to APAP challenge (Fig. 6A). Treatment restored circulating E2 levels (Fig. 6B) and reversed uterine atrophy (Fig. 6C), confirming effective hormone replacement. Following APAP overdose, E_2_ supplementation largely reversed the hepatoprotective phenotype of OVX females, resulting in significantly increased hepatocellular death compared with untreated OVX controls (Fig. 6D).

**Figure 6.**
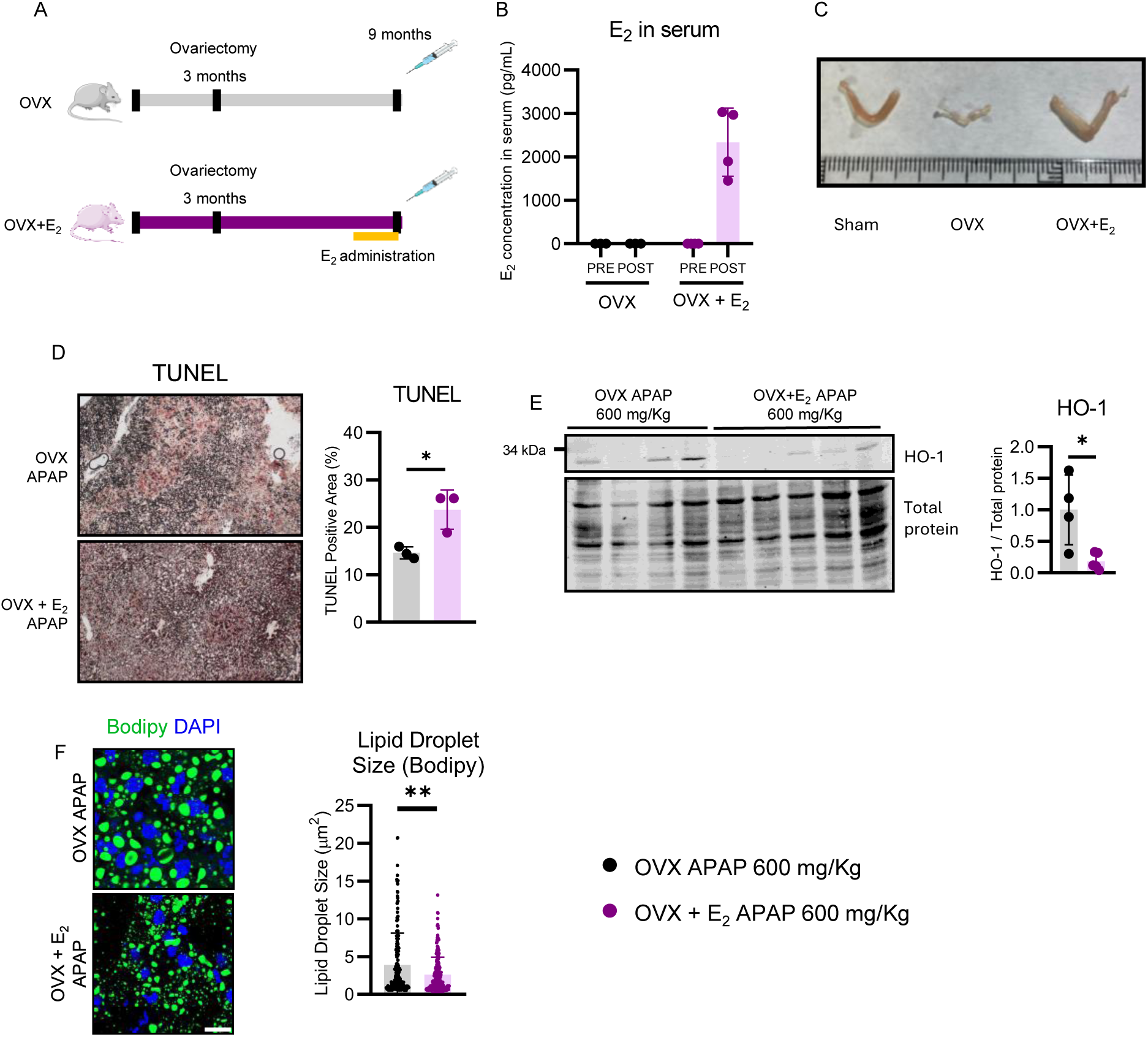
Estradiol supplementation increases susceptibility to APAP-induced liver injury and promotes lipid droplet expansion in OVX mice. (A) Schematic representation of the experimental design and study workflow. (B) Serum estradiol (E2) concentrations in the indicated experimental groups. (C) Representative images of uterine morphology from the indicated groups. (D) Representative TUNEL staining of liver sections and quantification of TUNEL-positive necrotic areas (n = 3-4 animals per group). Brown staining indicates necrotic tissue, whereas green staining indicates viable tissue (scale bar = 500 μm). (E) Representative immunoblot of HO-1 in whole liver lysates and corresponding densitometric quantification normalized to total protein content, with each dot representing an individual animal (n = 4-5 animals per group). (F) Representative BODIPY staining of neutral lipids in liver sections and corresponding quantification of lipid droplet size (scale bar corresponds to 20 μm). 15 randomly selected lipid droplets per animal were quantified from 3-4 animals per group. Data are presented as mean ± SD. Statistical significance was assessed using unpaired Student’s t-tests. *p < 0.05, **p < 0.01, ***p < 0.001.

This loss of protection was accompanied by reduced NRF2 pathway activation, as evidenced by lower hepatic levels of the NRF2 target protein HO-1 (Fig. 6E). Interestingly, and consistent with our previous observations, E_2_ supplementation also led to a marked reduction in hepatic lipid droplet size compared with untreated OVX mice (Fig. 6F).

These findings indicate that estradiol replacement reverses the lipid-droplet remodeling and hepatoprotective phenotype induced by long-term estrogen deprivation. E2 supplementation reduced NRF2 activation while increasing hepatocellular injury, supporting estradiol as an upstream regulator of the adaptive metabolic and antioxidant response to APAP. Together, these data link hormonal status, hepatic lipid remodeling, and NRF2-dependent cytoprotection.

### An age-associated hepatic signature in women links NRF2 activation with lipid metabolic remodeling

Finally, to assess the translational relevance of our findings, we analyzed hepatic mRNA expression of *NRF2* and its downstream targets *HO-1* and *SQSTM1* in human liver samples without histological evidence of steatosis. Samples were stratified into women <50 years, women ≥50 years, and men, with age ≥50 years used as a proxy for menopausal status [27]. Women aged 50 years or older exhibited increased hepatic expression of NRF2 and its downstream targets *HO-1* and *SQSTM1*, together with elevated expression of lipid metabolic genes including *FASN*, *ACC*, and *PPARA*, consistent with coordinated activation of NRF2 signaling and lipid remodeling programs (Fig. 7A-F). Collectively, these findings identify an age-associated hepatic transcriptional signature in women characterized by enhanced NRF2 signaling together with altered lipid metabolic gene expression. This molecular profile recapitulates key features observed in OVX mice, supporting the translational relevance of the experimental model. Although menopausal status was not directly available in this cohort, the concordance between the human and experimental data suggests that long-term estrogen deprivation may contribute to conserved metabolic and antioxidant adaptations in the liver.

**Figure 7.**
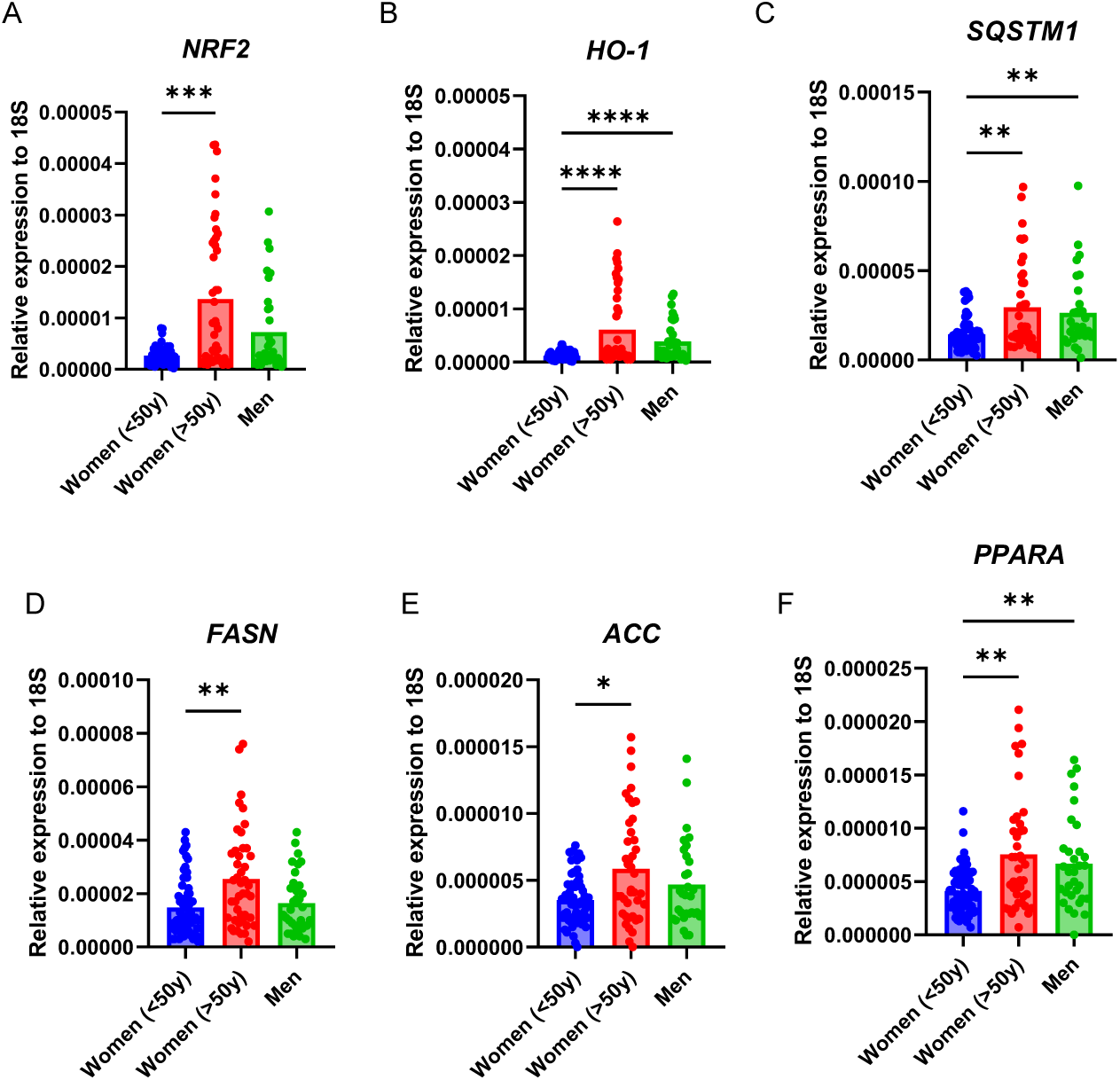
Postmenopausal women display increased expression of NRF2-responsive genes and genes involved in lipid synthesis. (A-F) Relative hepatic mRNA expression of the indicated genes in women <50 years (n=60), women >50 years (n=44), and men (n=33), normalized to 18S expression. Each dot represents an individual subject. Statistical analyses were performed using the Kruskal–Wallis test, followed by Dunn’s multiple-comparisons test. A p value < 0.05 was considered statistically significant.

## DISCUSSION

Menopause represents a critical stage in women’s health, characterized by hormonal changes that increase the risk of cardiovascular and metabolic diseases [6,7,27,5,3]. While the hepatic impact of menopause has been predominantly studied in the context of metabolic dysfunction and steatotic liver disease [1], the potential influence on acute liver injury remains poorly understood.

Despite increasing evidence that sex and age influence susceptibility to AP-ILI [12,13] little is known about how the adaptive changes that develop during long-term estrogen deprivation shape the hepatic response to APAP toxicity.

Here, we provide evidence that postmenopausal women exhibit a lower susceptibility to severe AP-ILI and that female mice subjected to long-term estrogen deprivation display increased resistance to APAP hepatotoxicity. This protective phenotype is associated with profound hepatic lipid remodeling, sustained activation of the NRF2 pathway, and an enhanced antioxidant response.

Importantly, NRF2 silencing attenuated the protective phenotype and increased susceptibility to APAP-induced injury, supporting a major contribution of this pathway to hepatic resilience following estrogen deprivation, although additional mechanisms are also likely involved.

A major finding of our study is that long-term estrogen deprivation induces extensive hepatic lipid remodeling characterized not simply by lipid accumulation, but by a shift toward a more saturated lipid environment and enlarged lipid droplets. These adaptations may represent a coordinated mechanism that enables hepatocytes to accommodate the metabolic challenges associated with estrogen withdrawal. This interpretation is consistent with previous studies showing that unsaturated fatty acids promote triglyceride synthesis and lipid droplet formation, thereby limiting lipotoxicity [19], whereas saturated phospholipids favor the formation of larger lipid droplets [29]. Accordingly, the lipid phenotype observed in OVX mice may reflect a state of metabolic adaptation that enhances tolerance to cellular stress. NRF2 activation appears to be a key component of this adaptive response. As a central regulator of cellular defense against oxidative, xenobiotic, and lipotoxic stress [20,30], NRF2 has been increasingly linked to lipid metabolism and lipid droplet dynamics. Consistent with this concept, short-term high-fat diet feeding reproduced both NRF2 activation and the hepatoprotective phenotype observed in OVX mice. Moreover, following APAP challenge, OVX livers were enriched in antioxidant lipids, including plasmalogens and CoQ9 [31–34], and plasmalogen metabolism has been reported to interact with NRF2 signaling [35]. Together, these findings support a model in which lipid remodeling contributes to sustained NRF2 activation and hepatic preconditioning during long-term estrogen deprivation, although the temporal and mechanistic relationship between these processes remains to be established. Similar protective responses have also been reported in MASLD models of APAP-induced liver injury involving PPARγ, a key regulator of lipid droplet biogenesis [36]. Estradiol supplementation reversed the hepatoprotective phenotype of OVX mice, restoring APAP susceptibility, suppressing NRF2 signaling, and normalizing hepatic lipid droplet architecture. This identifies estradiol as a key regulator of the adaptive program induced by long-term estrogen deprivation. Given the widespread use of hormone replacement therapies in postmenopausal women, these observations may have implications for the interpretation of individual responses to hepatotoxic insults.

The translational relevance of our findings is strengthened by human data showing that women older than 50 years exhibit a lower susceptibility to AP-ILI and increased hepatic expression of NRF2-responsive and lipid metabolism genes. These observations are consistent with reports from the Drug-Induced Liver Injury Network (DILIN), which identified greater severity of liver injury in women younger than 50 years compared with men and older women [37]. While age is an imperfect surrogate for menopausal status and does not establish causality, the concordance between the human data and the OVX model supports the clinical relevance of the adaptive pathways identified here. These findings suggest that endocrine aging may influence susceptibility to AP-ILI. Given that hormonal status is rarely considered in drug dosing or risk stratification, our results warrant further investigation into its impact on hepatic responses to pharmacological agents.

Although identified in the context of APAP hepatotoxicity, metabolic adaptations due to the long-term estrogen deprivation may have broader implications for hepatic responses to other xenobiotics. This is particularly relevant for postmenopausal women, a growing population frequently exposed to chronic pharmacological treatments. Whether these mechanisms are APAP-specific or represent a more general hepatoprotective adaptation remains to be determined.

Beyond xenobiotic response, our findings suggest that menopause-associated hepatic lipid remodeling is not inherently deleterious but may initially represent an adaptive response aimed at preserving hepatic homeostasis. We speculate that estrogen loss constitutes a first metabolic challenge that activates compensatory mechanisms, including NRF2 signaling, whereas progression to overt pathology requires additional insults that exceed this adaptive capacity. In this context, maintenance of hepatic homeostasis may depend on a delicate balance between sufficient NRF2 activity to promote stress resilience and avoidance of chronic or excessive pathway activation. Understanding the mechanisms that regulate this transition from adaptation to pathology may provide new opportunities for therapeutic intervention during menopause.

## MATERIAL AND METHODS

### Human studies

Clinical cohort: DILI risk was evaluated in a multicenter cohort of 5,112 patients admitted for paracetamol overdose (2008-2024). Patients aged 35-49 years were excluded, and the association between sex/age groups and severe hepatotoxicity (peak ALT >1000 U/L) was assessed using Firth-penalized logistic regression adjusted for age, admission ALT, and paracetamol levels. Analyses were approved by Edinburgh Medical School Research Ethics Committee (ref. 25-EMREC-070).

Human liver biopsies: Liver samples from obese individuals undergoing bariatric surgery were collected to quantify hepatic gene expression by RT-qPCR. Total RNA was isolated, reverse-transcribed, and analyzed by SYBR Green qPCR using 18S rRNA as the reference gene. All studies were conducted with ethical approval and informed consent. The study was performed in agreement with the Declaration of Helsinki according to local and national laws and was approved by the Ethics and Clinical Research Committee of the Virgen de la Arrixaca University Hospital (ref. number 2020-2-4-HCUVA) and written informed consent was obtained from each participant.

### Animal model

All animal procedures were performed according to the EU Directive 86/609 EEC and Recommendation 2007/526EC and were approved by the Ethical Comitee from Autonomous University of Madrid and CICbioGUNE Animal Care and Use Committee and the local authority (Comunidad de Madrid and Diputación de Bizkaia) according to the criteria established by the European Union. Female C57BL/6J mice underwent bilateral ovariectomy or sham surgery at 12 weeks of age and were subsequently challenged with APAP overdose (600 or 360 mg/kg, i.p.) under different nutritional conditions; in selected experiments, OVX mice received daily estradiol supplementation for 15 days before APAP administration. NRF2 was silenced in vivo using a predesigned siRNA formulated with Invivofectamine™ 3.0 and administered by intravenous tail-vein injection; 24 h later, mice were challenged with a high-dose APAP overdose as described above.

## Conflict of interest statement

The authors declare no conflicts of interest.

## Financial support statement

Research in the S.C. lab is supported by the Grant PID2020-114054RA-I00 and PID2023-148516OB-I00 founded by MCIN/AEI/10.13039/501100011033 and CNS2023-143646 funded by MICIU/AEI/10.13039/501100011033, the European Union NextGenerationEU/PRTR. J.G-C is supported by the fellowship PRE2021-097661 founded by MCIN/AEI/10.13039/501100011033.

J.M.L-A is supported by the fellowship PRE2022-000715 founded by PID2022-136738OB-I00. A Q.G.M. is supported by the fellowship PRE2022-105893.

This work was supported by grants from Ministerio de Ciencia, Innovación y Universidades MICINN: PID2023-146933OB-100 funded by MCIU /AEI /10.13039/501100011033 / FEDER, UE as part of Plan Estatal de Investigación Científica y Técnica e Innovación (to MLM-C); Grant CEX2021-001136-S funded by MCIN/AEI /10.13039/501100011033 (to MLM-C). This work was supported by the Spanish Ministry of Science, Innovation and Universities (PID2024-161269OB-I00 to EA-G).

This work was supported by grants to PG: AIRC (MFAG 2020), PNRR (PNRR-MR1-2022-12376821), PNRR (PNRR-MCNT2-2023-12377641), PRIN-2022-PNRR (P2022JLNZ7), PRIN-2022 (20224FL9T5), Fondazione Telethon.

This work was supported by the Fundación Séneca (Grant number 22508/PI/24), and by the Institute of Health Carlos III (ISCIII) (Grant number PI23/00171); co-funded by the European Union).

## Authors contributions

José García-Consuegra: Investigation, Writing – Original Draft.

Naroa Goikoetxea-Usandizaga: Investigation, Writing – Original Draft.

Ariadna Qin Gil Martín: Investigation.

Ana Sagrera Aparisi: Investigation.

José María Lanza-Arnaiz: Investigation.

Rene González-Recio: Investigation.

Carolina Conter: Investigation.

Raúl J. Andrade: Resources.

M. Isabel Lucena: Resources.

Paolo Grumati: Investigation.

Bruno Ramos-Molina: Investigation, Resources.

Estela Area: Investigation.

James W. Dear: Investigation, Formal Analysis, Resources.

Chris Humphries: Investigation, Formal Analysis, Resources.

Laura Formentini: Resources.

María Luz Martínez-Chantar: Conceptualization, Resources, Writing – Original Draft, Funding Acquisition.

Sara Cogliati: Conceptualization, Resources, Writing – Original Draft, Funding Acquisition.

## Data availability statement

Human electronic health record data from Edinburgh are not available due to confidentiality restrictions on onward sharing.

## Acknowledgment

We acknowledge the Advanced Light Microscopy Facility (SMOA), the Electron Microscopy Facility (SME), the Animal Facility and the Biocomputational and Genomics Facilities at the Centro de Biologia Molecular Severo Ochoa (CBM); the Mass Spectrometry Facility at TIGEM.

## SUPPLEMENTARY FIGURES LEGENDS

**Figure S1.**
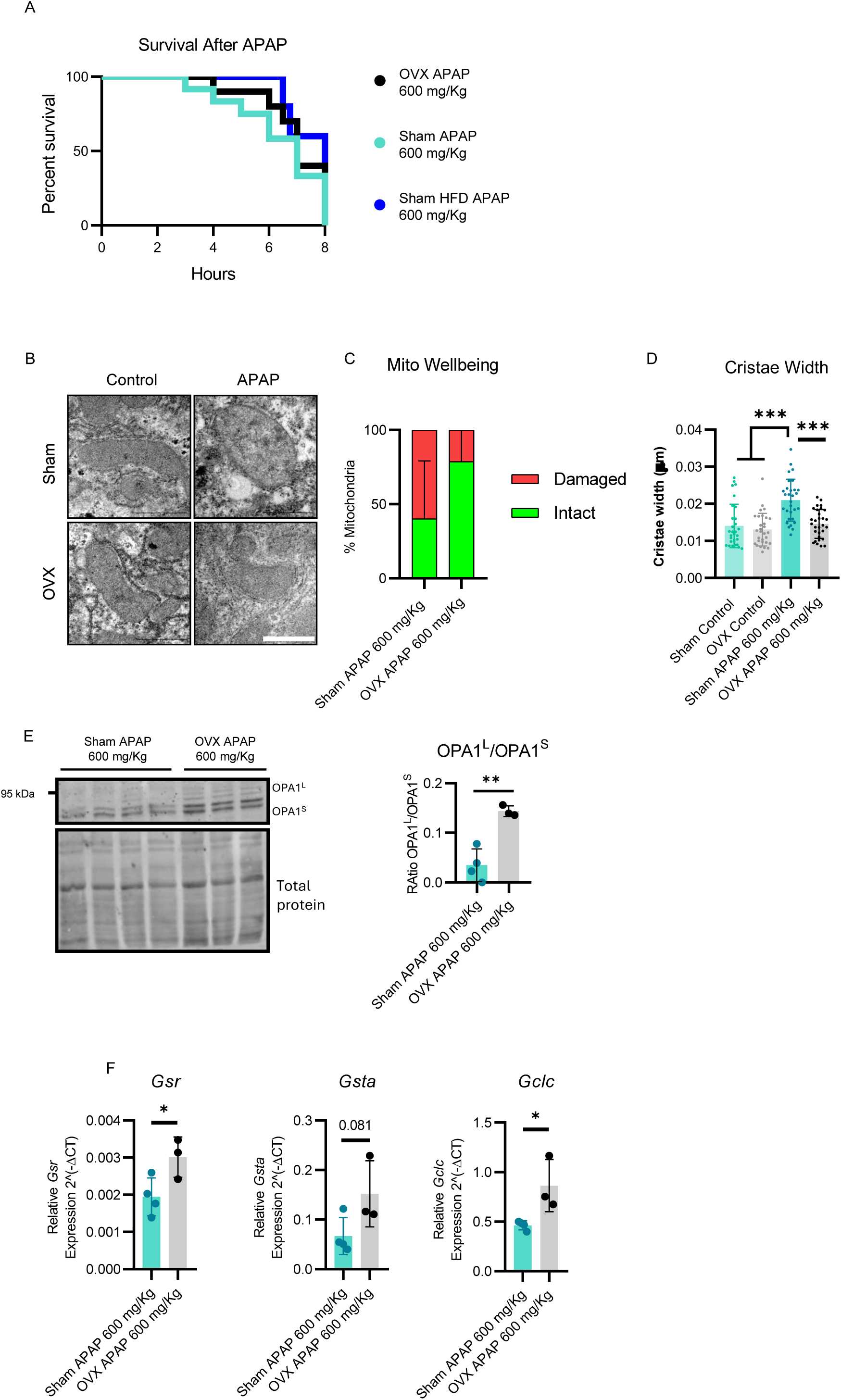
Ovariectomy improves survival following APAP overdose and alters mitochondrial morphology. (A) Kaplan-Meier survival curves of the indicated experimental groups following APAP overdose. (B) Representative transmission electron microscopy (TEM) images of liver sections from the indicated experimental groups (scale bar corresponds to 1 μm). (C) Quantification of mitochondrial wellbeing based on TEM analyses (n = 3 animals). (D) Quantification of mitochondrial cristae width. (E) Representative immunoblots of OPA1 in whole liver lysates and corresponding densitometric quantification of the ratio between long OPA1 isoforms and short soluble OPA1 forms. (F) Relative hepatic mRNA expression of *Gsr, Gsta and Gclc* normalized to Arp expression (n = 3-4 animals per group). Data are presented as mean ± SD. One Way ANOVA was used to analyze (D), while (E) and (F) was analyzed with unpaired Student’s t-tests. **p < 0.01, ***p < 0.001.

**Figure S2.**
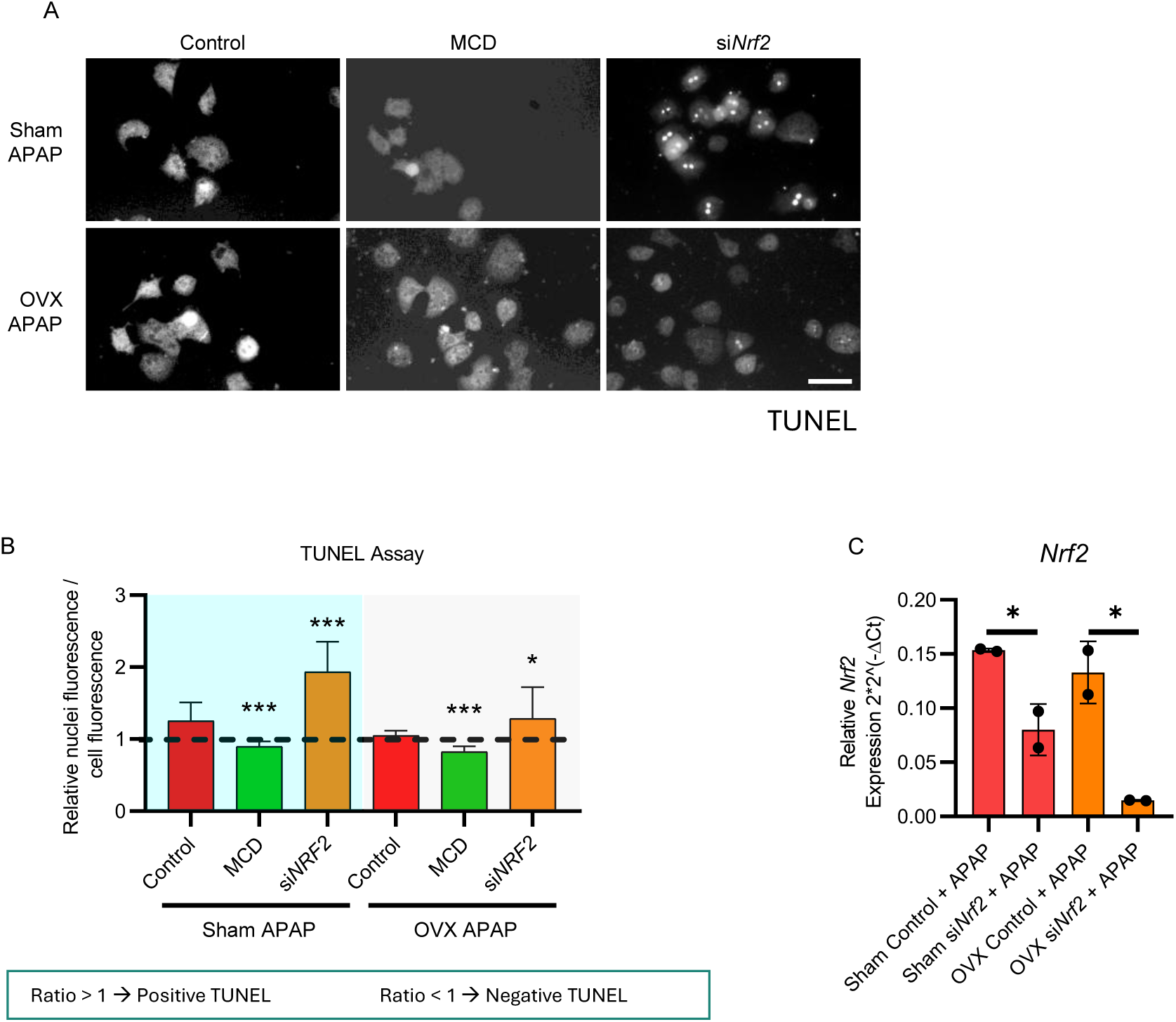
*Nrf2* silencing and culture medium composition modulate APAP-induced cell death in primary hepatocytes. (A) Representative TUNEL staining of primary hepatocytes isolated from the indicated experimental groups and cultured under the specified conditions, including Nrf2 silencing (*siNrf2*) and methionine- and choline-deficient medium (MCD) followed by APAP treatment. Corresponding quantification of TUNEL-positive cells is shown (scale bar = 50 μm). (B) TUNEL positivity was determined as the ratio between nuclear TUNEL fluorescence intensity and background signal intensity, with values > 1 considered positive. Each dot represents an individual cell (n = 18-39 cells from 3 technical replicate). (C) Relative mRNA expression of *Nrf2*, normalized to *Arp* expression (n=2 technical replicates). Data are presented as mean ± SD. Statistical significance was assessed using unpaired Student’s t-tests comparing each condition vs its control *p < 0.05.**p < 0.01, ***p < 0.001.

**Figure S3.**
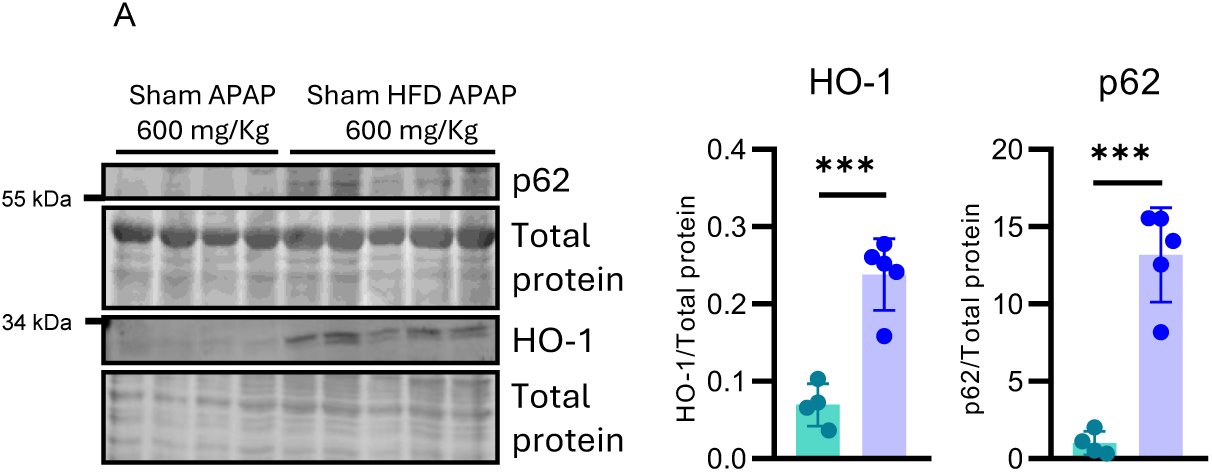
HFD induces NRF2 activation following APAP overdose. **(A)** Representative immunoblots of HO-1 and p62 in whole liver lysates and corresponding densitometric quantification normalized to total protein content, with each dot representing an individual animal (n = 4-5 animals per group). Data are presented as mean ± SD. Statistical significance was assessed using unpaired Student’s t-tests. ***p < 0.001.

## SUPPLEMENTARY MATERIALS

### Human studies

We assessed the DILI risk in a multi-site (3 hospital) cohort of all patients admitted to NHS Lothian hospitals in Edinburgh, UK, between 2008 and 2024, who received an ICD-10 code of T39.1 (Poisoning by 4-Aminophenol derivatives). Patients were included if they had a paracetamol level and an ALT level measured within a clinical window of 24 hours prior to 4 hours after the time of admission (i.e. obtained within the Emergency Department prior to hospital ward admission, or shortly following hospital ward admission). If paracetamol values were below the limit of detection (<5mg/L) they were imputed as 2.5mg/L. Modelling was restricted to admissions with a presentation ALT <1000. To maintain clear biological separation between the groups and minimize the influence of the perimenopausal transition, all patients aged 35 to 49 years were excluded from the primary analysis. This resulted in a final cohort of 5,112 admissions. To determine the independent association between cohorts and the risk of hepatotoxicity (peak ALT >1000 U/L), we performed a Firth-penalized logistic regression. The model specifically compared Men<35, Men≥50, and Women≥50 against the reference group (Women<35). To isolate the cohort effect while accounting for residual confounding, we adjusted for continuous age, log-transformed presentation ALT, and log-transformed presentation paracetamol levels. Adjusted odds ratios (aOR) and 95% confidence intervals were calculated, with an aOR < 1 indicating a lower likelihood of severe injury relative to the reference group. Statistical significance was defined as p< 0.05, without multiplicity correction due to the exploratory nature of analyses.

### RT-qPCR of human liver biopsies

A total of 137 patients with obesity who underwent bariatric surgery at the Virgen de la Arrixaca University Hospital (Murcia, Spain) between January 2020 and December 2022 were included in the study. Patients were aged 18-65 years and provided written informed consent, age 18–65 years. Liver biopsies of at least 1 cm2 were collected during surgery. Total RNA was extracted from 10–20 mg of liver tissue using TRIzol-chloroform protocol. RNA was eluted in RNAse-free water, and its concentration and purity were assessed using a Nanodrop spectrophotometer. Only samples with an A260/A280 ratio between 1.8 and 2.1 were used for gene expression analyses.

A total of 1 μg of RNA was reverse-transcribed using the High-Capacity RNA-to-cDNA kit (Applied Biosystems), following the manufacturer’s instructions. Negative controls without RNA template and controls without reverse transcriptase were included to assess potential contamination and genomic DNA amplification, respectively.

Real-time quantitative PCR (RT-qPCR) was carried out using Power SYBR Green Master mix (Applied Biosystems, Thermo Fisher Scientific) on a Fast 7500 Real-Time PCR System (Applied Biosystems, Thermo Fisher Scientific). Relative gene expression was calculated using the 2^−ΔCT method, with 18S rRNA as the reference gene. The primers sequences used are listed below.

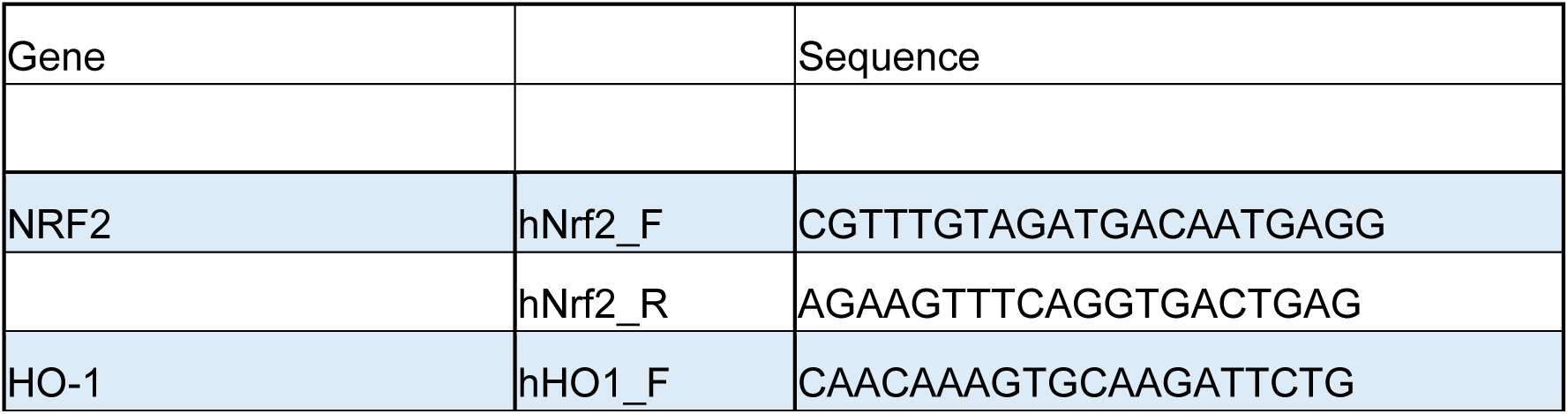
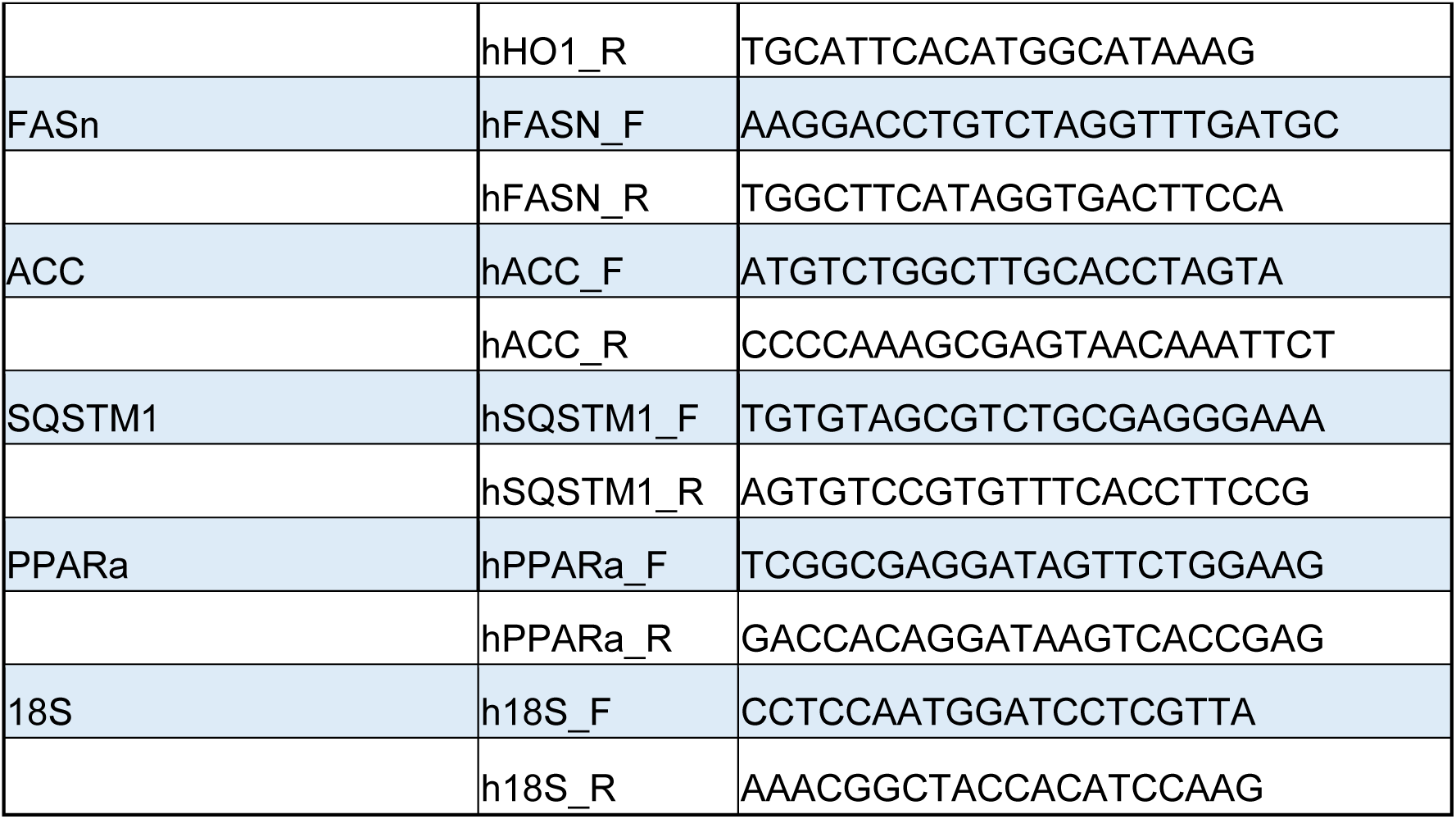

The study was performed in agreement with the Declaration of Helsinki according to local and national laws and was approved by the Ethics and Clinical Research Committee of the Virgen de la Arrixaca University Hospital (ref. number 2020-2-4-HCUVA) and written informed consent was obtained from each participant.

### Animal model

All animal procedures were performed according to the EU Directive 2010/63/UE and were approved by the Comité de Ética from Autonomous University of Madrid. Animal procedures were approved by the CICbioGUNE Animal Care and Use Committee and the local authority (Diputación de Bizkaia) according to the criteria established by the European Union.C57BL/6JOlaHsd mice were housed in the Centro de Biología Molecular Severo Ochoa (Madrid) and CICBioGUNE (Bilbao) facilities, with a constant temperature of 21°C and 12/12 hours light/dark cycles, and fed with water and a standard chow diet ad libitum except when specified.

### Ovariectomy

Bilateral ovariectomy was performed at 12-weeks old female mice. In brief, animals were anesthetized with isoflurane (5% induction, 2% for maintaining) and injected with Rimadyl (572101, LEONVET, 10 mg/Kg) and Buprenorphine (587424, LEONVET; 0,1 mg/Kg) intraperitoneally. Both sides of the animal’s back (skin and muscle) were cut, and both ovaries were removed by using a cauteriser. To close the incision, the muscle was closed with surgical suture, while the skin was closed with surgical grips. Sham animals underwent the same procedure, except the ovary removal and ovariectomy was further confirmed by the analysis of the vaginal wet smear and uterus size at the time of sacrifice.

### In vivo acetaminophen (APAP) overdose

Mice were starved for 12 h and received a single dose of acetaminophen (APAP, Sigma-Aldrich, A7085-100g) 600 mg/kg intraperitoneally. All animals were monitored every 15 minutes. Tissues were collected immediately upon spontaneous death or, if the animals survived, at 8 hours after APAP injection, when all remaining animals were euthanized and the tissues were collected for subsequent analyses.

Where indicated, animals were fed with Chow Diet or High Fat Diet (HFD,60% fat,Research Diets, D12492i) 5 days prior APAP injection. Caloric restriction protocol was applied were indicated and, in that case, animals were fed with a food amount equivalent to 40% of their food intake, previously registered, for 5 consecutive days prior the APAP overdose. To perform the in vitro assays with primary hepatocytes and the measurements of transaminases, mice were starved for 12 h and injected with a single dose of 360 mg/Kg. 24 hours after, the isolation and culture of primary hepatocytes and blood extraction was performed.

### Estradiol in vivo supplementation

For the estradiol treatment, 1.25 μg of pure estradiol (Sigma, E4389) was dissolved in 150 µL of 0.9% saline solution (NaCl). Mice received daily intraperitoneal injections of either 150 μL of the estradiol solution or 150 μL of the vehicle for 15 consecutive days prior to APAP administration.

### Estradiol detection kit

The 17-β Estradiol Elisa kit (Abcam, ab108667) was used to further confirm the success of the exogenous supplementation of the hormone by following manufacturer’s instructions.

### Subcellular fractionation

30 mg of liver tissue were homogenized in medium A (320 mM sucrose, 10 mM Tris-HCl, 1 mM EDTA, pH 7.4) with a potter-Elvehjem associated with a Hei-TORQUE Expert 100 mechanical shaker (Heidolph) with 5-6 strokes at 600 rpm. Homogenates were further centrifuged for 10 minutes at 4°C at 1000 xg. While the surnatant was considered as the “cytosolic fraction”, the pellets were kept as “nuclear fraction”. The protein concentrationwere then quantified with BCA and prepared as previously described for SDS-PAGE. To validate the success of the subcellular fractionation, nuclear and cytosolic markers (antibodies anti H3 and GAPDH) were used.

### Quantification of GSSG GSH and GSH+NAPQ1 metabolites

Relative amount of the indicated metabolites was determined with a UPLC system (Acquity Waters, Manchester) coupled to a Time-of-Flight mass spectrometer (ToF MS, SYNAPT G2, Waters). A 2.1 × 100 mm, 1.7 mm BEH amide column (Waters), stabilized at 40 °C, was used to separate the analytes before entering the MS. The aqueous phase consisted of 99.5% water, 0.5% formic acid, and 20 mM ammonium formate while the organic phase consisted of 29.5% water, 70% MeCN, 0.5% formic acid, and 1 mM ammonium formate. The extracted ion trace was obtained for GSH (m/z = 308.0916), GSSG (m/z = 613.1598) and NAPQ1-GSH (m/z = 457.139) in a 20 mDa window and subsequently smoothed and integrated with TargetLynx software (Waters, Manchester).

### Blood collection

Blood samples were collected by retro-orbital puncture using heparinized capillary tubes and transferred to serum separator gel tubes (Microtainer, Becton Dickinson). Samples were centrifuged at 2,500 × g for 5 minutes at 4°C to obtain serum. Serum alanine aminotransferase (ALT) and aspartate aminotransferase (AST) activities were measured using a Selectra Junior Spinlab 100 analyzer (Vital Scientific).

### Primary hepatocyte isolation

Primary hepatocytes were isolated from the indicated mice 24 hours after the injection with 360 mg/kg of APAP. Briefly, mice were anesthetized with isoflurane (4% in O2 for induction and 1,5% for maintenance). The mouse’s abdomen was opened, and a catheter was inserted into the inferior cava vein. Liver was perfused with Buffer I (Glucose 22mM, NaCl 122 mM, KCl 4,9 mM, NaHCO3 25 mM, KH2PO4 1,2 mM, MgSO4 2,9 mM, 0,31mM EGTA), Buffer II (same as Buffer I without EGTA) and Buffer III (same as Buffer 1 + Collagenase Type IV, 1mg/mL). After, the liver was placed in a petri dish and was mechanically disrupted in Minimum essential medium (MEM; Gibco) supplemented with 1% penicillin (100 U/mL), streptomycin (100 u/mL), and glutamine (1%). The homogenate was then filtered through a sterile gauze and washed three times in MEM (2x 48 xg 5 minutes at 4°C and a last wash at 58 xg for 5 minutes at the same temperature). Hepatocytes were then seeded in collagen-coated tissue culture dishes in 10%FBS MEM + 1% PS and 1% Glutamine and maintained in a 5% CO2-95% air incubator during 6h. After that, the medium was retired and replaced by 5% FBS MEM supplemented with 1% PS and 1% Glutamine. 24 hours after the cell seeding, the experiments were performed.

### In vitro APAP treatment

The in vitro APAP treatment for primary hepatocytes started following the standard protocol to isolate primary cells from WT and OVX mice. The cells were seeded and let for 6 hours before changing the medium to fresh supplemented MEM, methionince-coline deficient medium (MCD) or Krebs-Henseleit-HEPES (KHH) medium (NaCl 120 mM, 4,7 mM, CaCl2 2,5 mM, MgSO4 1,2 mM, KH2PO41,2 mM, NaHCO3 25 mM, HEPES 25 mM pH 7.4) and let for 12 hours. After 12 hours, APAP was added to the primary hepatocytes at a concentration of 3 mg/mL for 6 hours. The cells were then washed with PBS and processed for the experiments.

### In vivo NRF2 silencing

NRF2 was silenced by using the Invivofectamine™ 3.0 Reagent (ThermoFisher Scientific, IVF3001) and a predesigned siRNA (ThermoFisher Scientific, 4457308, sequence 5’-3’:GCAUGUUACGUGAUGAGGAtt). Briefly, the siRNA was prepared as recommended by the Invivofectamine supplied protocol and the silencing mix was administered by intravenous tail injection. 24 hours after the injection, the animals were then administered with the high dose of APAP as previously described.

### In vitro NRF2 silencing

For the in vitro silencing of NRF2, hepatocytes were extracted and cultured in MEM 0%? FBS that was supplemented with the silencing mix, containing 0,50 nM of siRNA (Sigma-Merck, 00130750, sequence 5’-3’: GAAUUACAGUGUCUUAAUA) with DharmaFect (Dharmacon, T-2001-01). This mix was maintained for 6 hours, when the media was replaced with fresh MEM 5% FBS, which was let for 12 hours. After that, in vitro APAP administration was done as previously described.

### TUNEL Assay

TUNEL staining was performed in primary hepatocytes using the In Situ Cell Death Detection Kit (Roche, 11684795910) according to the manufacturer’s instructions. For liver sections, the TUNEL Assay Kit (Abcam, ab206386) was used following the manufacturer’s protocol. Stained liver sections were imaged using a DMC6200 CMOS Color camera, whereas primary hepatocytes were imaged with either a Leica SP8 Lightning confocal microscope or an Axio Imager D1 upright fluorescence microscope.

### Triglycerides measurement

To measure the triglycerides’ levels within the hepatic tissue, the Triglyceride Assay Kit (Abcam, ab65336 was used following the manufacturer’s instructions.

### Bodipy Staining

Bodipy staining was done in fixed liver slices. Briefly, the liver was frozen in OCT and further cut in cryosections. These slices were let defrost and washed twice with PBS prior the staining with a mix containing Bodipy 2 µM, 0,5% Ethanol in PBS. The mix was let in dark for 1 hour. After, the mix was retired and two washes with PBS were done prior the staining of the nuclei with DAPI staining (1 µg/mL) for 10 minutes. Finally the sections were mounted with Prolong and visualized by using an Zeiss LSM800 inverted microscope.

### Respiration assay in primary hepatocytes

To assess the respiration of primary hepatocytes, a Seahorse Bioscience XF24 Extracellular Flux Analyzer was used. 24 hours after seeding 15.000 isolated primary hepatocytes in a collagen-coated XF24 cell culture microplate (Seahorse Bioscience), the medium was replaced with 500 µL of MEM containing 1mM sodium pyruvate, 2 mM L-glutamine and maintained at 37°C in a chamber without CO2. Oxygen consumption rate was determined upon the addition of oligomycin (1 mM), FCCP (300 nM) and Rotenone/Antimycin (4/6 µM). The normalization of the OCR was done using the total protein amount, measured with BCA reagent.

### Protein isolation

30 mg of liver tissue were lysated in RIPA buffer and incubated in ice for 20 minutes. The samples were then centrifuged for 20 minutes at 12.000 xg, 4°C to collect the supernatant. For primary cells, 1 confluent well of a MW6 plate was harvested in 100 µL of RIPA buffer and incubated in ice for 20 minutes. The enriched protein fractions were quantified with BCA reagent and finally stored at -80°C.

### SDS-PAGE and Western Blot

20 µg of isolated protein were boiled with 5X loading buffer (312.5 mM Tris-HCl, pH 6.8, 10% SDS, 50% glycerol, 0.05% bromophenol blue, and 25% β-mercaptoethanol), separated by SDS-PAGE and transferred onto polyvinylidene fluoride (PVDF) or nitrocellulose membranes. Primary antibodies were incubated at the indicated concentration for 2 hours at RT or O/N at 4°C (anti-HO-1 1:1000 SIGMA, AB1284; anti-p62/SQSTM1 (D6M5X) 1:1000, Cell Signaling Technology, 23214; Keap1 1:1000, ProteinTech, 10503-2-APM; anti-NRF2 1:500, Santa Cruz BiotechnologiesSC-722; anti-GAPDH, 1:1000 abcam, ab9485; anti-H3, 1:1000 abcam, ab18521; anti-OPA1, 1:1000 BD BioSicence, 612607) while the secondary antibodies were incubated for 1 hour at RT at a concentration of 1:5000 (anti-Mouse 800 Goat anti-Mouse fluorescent antibody, Invitrogen, SA5-10176; anti-Rabbit 680 Goat anti-Rabbit fluorescent antibody, Invitrogen, A21076).

Blots were then revealed using the Odyssey M (LI-COR) and all the analysis were performed using the ImageJ software. To normalize the signal, the membranes were stained with Staining Revert™ 520 Total Protein Stain (Licor) following the manufacturer’s protocol.

### Oil Red staining

Oil Red: Centro Nacional de Biotecnología (CNB-CSIC) histology facility performed the oil red staining procedure following standard protocols.

### Proteomics

All the experiments were performed in a labeling-free setting. Proteins were precipitated in acetone and peptides obtained using Preomics iST kit (https://www.preomics.com/). Instruments for LC MS/MS analysis consisted of a NanoLC 1200 coupled via a nano-electrospray ionization source to the quadrupole-based Q Exactive HF benchtop mass spectrometer. Peptides were separated based on their hydrophobicity using an analytical column (75 μm) in-house packed with C18-AQ resin (1.9 μm particles). The separation was performed with a linear gradient from 7% to 32% solvent B over 45 min, followed by 32% to 45% B in 5 min, 45% to 95% B in 3 min, and re-equilibration from 95% to 5% B in 5 min, at a flow rate of 300 nL/min. MS data were acquired in DIA (Data-Independent Acquisition) mode using 32 variable isolation windows covering a mass range of 300– 1650 m/z. The resolution was set to 60,000 for MS1 and 30,000 for MS2. The AGC target was 3 × 10⁶ for both MS1 and MS2, with maximum injection times of 60 ms and 54 ms, respectively. Normalized collision energies (NCE) were set to 25%, 27.5%, and 30%.

RAW files were analyzed using Spectronaut (v. 18, 19, 20), ran in library-free search (directDIA). To assess protein identification and quantification, default parameters were used: precursor ion mass tolerance: dynamic; Orbitrap fragment ion mass tolerance: dynamic (Spectronaut); fixed modifications: cysteine carbamidomethylation; variable modifications: methionine oxidation, protein N-terminus acetylation, enzyme: trypsin, Lys-C; the number of allowed missed cleavage sites: 2; minimum peptide length: seven amino acid residues. The DIA spectra were probed against UniProt *mus musculus* FASTA database. MS data were filtered at 1% FDR on the protein level and exported, followed by analysis with different software packages (Excel, Rstudio, SRplot). Specifically, statistical analyses for quantitative evaluation were performed using Perseus software (1.6.2.3). Subtraction of contaminants, reverse entries as well as the identification by a modified peptide only, were applied as the first filter. Missing values were replaced by random numbers that are drawn from a normal distribution. For the comparison among the different conditions in terms of protein abundance level, Anova and Student’s t test were used. For each test, the level of significance was set equal to 0.05. Proteins with at least log_2_ one fold changes were considered differentially regulated. A Gene Ontology (GO) and KEGG enrichment analysis was carried out on dysregulated proteins.

### Lipidomics

Lipids were extracted from equal amounts of material (30 mg protein/sample). Lipid extracts were prepared via chloroform–methanol extraction, spiked with appropriate internal standards, and analyzed using a Synap G2 LC/MS system (Waters coorporation). Cholesterol and cholesteryl esters were separated with normal-phase HPLC using an Agilent Zorbax Rx-Sil column (inner diameter 2.1 Å∼ 100 mm) under the following conditions: mobile phase A (chloroform:methanol:1 M ammonium hydroxide, 89.9:10:0.1, v/v/v) and mobile phase B (chloroform:methanol:water:ammonium hydroxide, 55:39.9:5:0.1, v/v/v/v); 95% A for 2 min, linear gradient to 30% A over 18 min and held for 3 min, and linear gradient to 95% A over 2 min and held for 6 min. Quantification of lipid species was accomplished using multiple reaction monitoring (MRM) transitions that were developed in earlier studies in conjunction with referencing of appropriate internal standards. Values are represented as mole fraction with respect to total lipid (% molarity). For this, lipid mass of any specific lipid is normalized by the total mass of all the lipids measured. Data distribution was assumed to be normal. Lipidomic data were normalized to the mean value of the reference group, which was set to 1. Data are presented as fold changes relative to this reference value and visualized using heatmaps.

### RNA isolation and RT-qPCR from mouse liver samples and primary hepatocytes

RNA was isolated following a standard trizol-chloroform protocol. Briefly, 30 mg of liver sample or 5*105 cells were homogenized in 1mL of Trizol (Invitrogen) and mixed with 200 µL of chloroform. The homogenate was then centrifuged at 12.000 xg for 15 minutes at 4°C and the supernatant was then passed to a new clean RNAase-free tube and mixed with 500 µL of isopropanol and finally stored at -80°C overnight. The day after, the RNA was pelleted after a centrifugation (12.000 xg, 15 minutes, 4°C). After two cleaning steps with pure EtOH, the pellet was finally resuspended in RNAase free water and quantified. 2 µg of total RNA were used as template to transcribe into cDNA using MLV reverse transcriptase. Quantitative real-time PCRs were done with Viia7 Real-Time PCR System (Applied Biosystems) or CFX OPUS 384 (Bio-Rad). The relative expression of mRNA levels were normalized with ARP mRNA levels as housekeeping gene. The amplification setup was 40 cycles in 384-well plates. The used primers were:

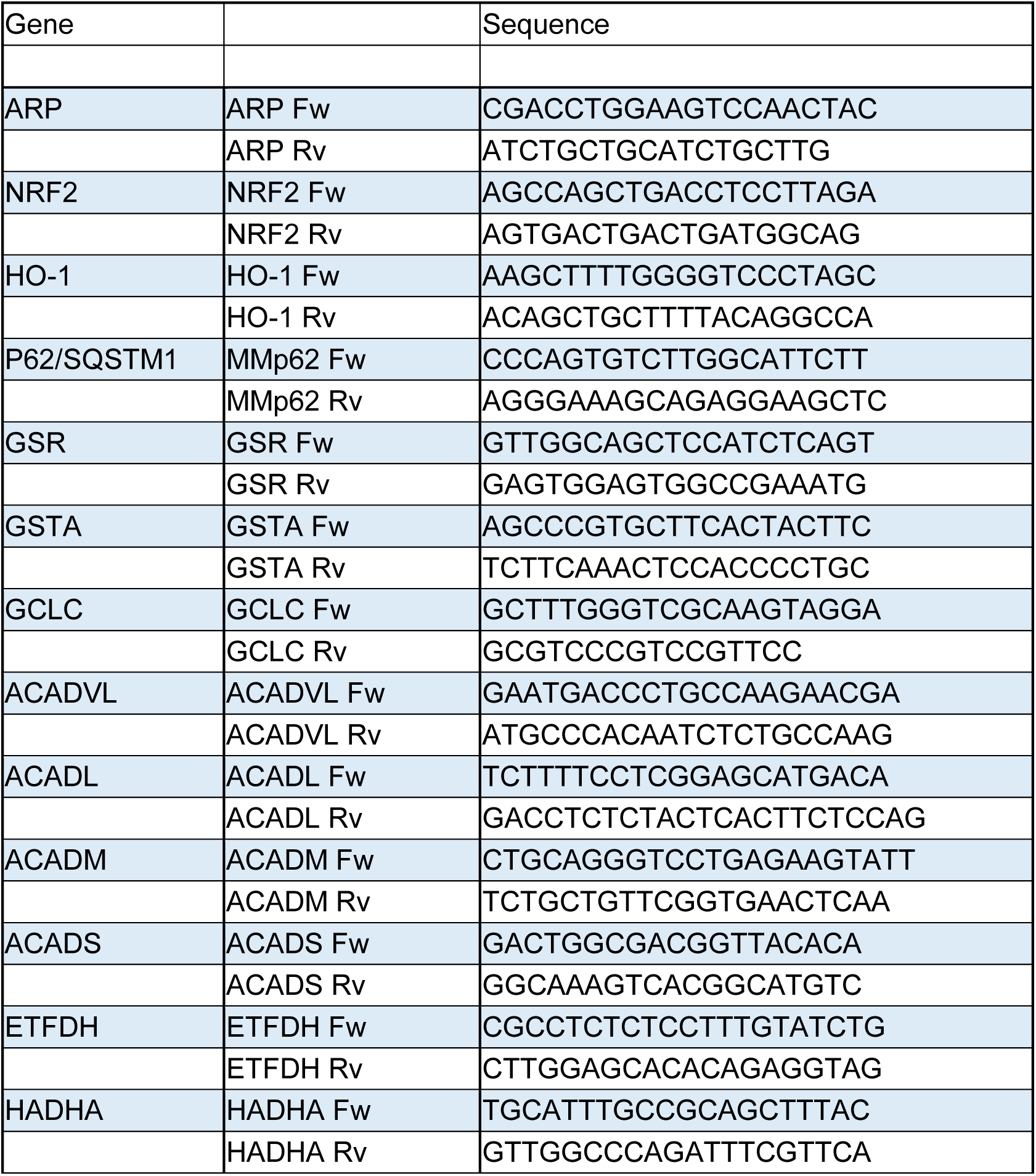
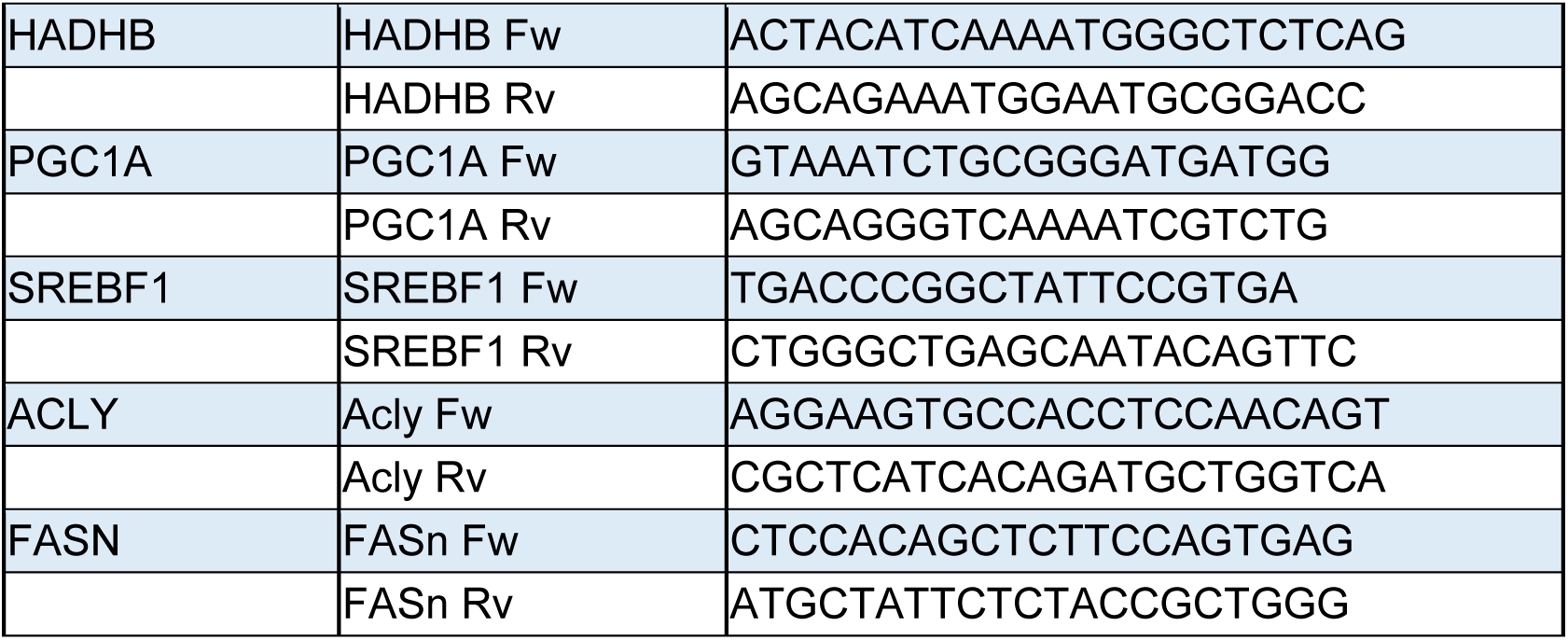

### Immunofluorescence

Primary hepatocytes were seeded in collagen-coated covers as previously described. Cells were fixed with 4% paraformaldehyde in PBS for 15 minutes, followed by two washes of PBS. The first incubation of NaBH4 was done to avoid autofluorescence (1 mg/mL in PBS) for 10 minutes RT, followed by 3 washes of PBS. Cells were then permeabilized with permeabilization buffer (Triton x100 0,2% in PBS) for 10 minutes and the blocking step was performed (1% BSA in permeabilization Buffer). Primary NRF2 antibody (Santa Cruz Biotechnologies, sc-722) was then incubated overnight in a wet chamber at a final dilution of 1:100.

The primary antibody was removed and 3 washes with PBS were done prior the incubation with the secondary antibody (Absorbed Secondary Antibody Alexa Fluor 555 conjugate, ThermoFisher A-31572) at a dilution 1:200 for 1 hour. 3 wash steps were done with PBS and the nuclei were counterstained with DAPI 1 μg/ml (268298, Merck) and the covers were mounted with Mowiol-Dabco (Advanced light microscopy Facility, CBMSO). Images were taken using an upright fluorescent microscope (Axioimager D1).

### Intracellular ATP Measurement

Hepatocytes were used to quantify the ATP production following the ATPlite luminescence ATP detection kit (PerkinElmer) following the manufacturer’s instruction. Briefly, cells were lysed using the cell lysis solution and further incubated on an orbital shaker (700 rpm, 5 minutes, RT). Substrate solution was then added and again incubated (700 rpm, 5 minutes, RT). After 10 minutes in dark, the luminescence was measured in a luminescence reader and ATP was extrapolated to an ATP standard curve and further normalized with the total protein content, measured by the BCA assay.

### Electron microscopy

Tissues were fixed with 4% paraformaldehyde and 2% glutaraldehyde in 0.1 M? phosphate buffer pH 7.4 overnight. Then, the tissue was embedded in agarose 6% and slices of 200 µm were cut in a Vibratome VT1200S (Leica). After, the sample’s preparation was followed by the Electron Microscopy Unit at the CBMSO. Briefly, the slices were treated with 1% osmium tetroxide at 4°C for 1 hour and then dehydrated in graded ethanol and embedded in TAAB 812 epoxy resin. Later, ultrathin serial sections (80 nm) of the sample were obtained using an ultramicrotome UltraCult E (Leica) and mounted on carbon coated cooper mesh grids. Sections were finally stained with 2% uranyl acetate for 7 minutes and lead citrate for 2 minutes and observed through a JEM 1400 Flash (Jeol) Transmission Electron Microscope and pictures were taken with a camera CMOS OneView (Gatan).

### Image Analysis

Image analysis was performed with ImageJ 1.54 software (Fiji).

### Statistics

Statistical analyses were done with GraphPad Prism 8.1 and Excel Software. All the performed analysis are indicated in the figure legend.

## ABBREVIATIONS

ACC: Acetyl-CoA Carbolxylase
AP-ILI: APAP-induced liver injury
ALF: Acute liver failure
aOR: adjusted odds ratio
APAP: Acetaminophen
CoQ9: Coenzyme Q9
Cyp2e1: Cytochrome P450 2e1
DILI: Drug-induced liver injury
DILIN: Drug-induced liver injury network
E2: 17β-Estradiol
FASN: Fatty Acid Synthase
GCLC: Glutamate-cysteine ligase catalytic subunit
GSH: Reduced Glutathione
GSR: Glutathione reductase
GSSG: Oxidized glutathione
GSTA: Glutathione S-transferase Alpha
HFD: High-fat diet
HO-1: Heme oxygenase-1
KEAP1: Kelch-like ECH-associated protein 1
MASLD: Metabolic dysfunction-associated steatotic liver disease
MCD: Methionine-and choline-deficient
NAPQI: N-acetyl-p-benzoquinone imine
NRF2: Nuclear factor erythroid 2-related factor 2
OPA1: Optic atrophy 1
OVX: Ovariectomy/ovariectomized
Peos: Plasmalogen phosphatidylethanolamine
PPARA: Peroxisome proliferator-activated receptor alpha
PUFA: Polyunsaturated fatty acid
SQSTM1: sequestosome-1 (p62)
SRXN1: sulfiredoxin-1
TAG: Triacylglycerol
TEM: Transmission electron microscopy
TUNEL: terminal deoxynucleotidyl transferase dUTP nick end labeling

## Notes

### Competing Interest Statement

The authors have declared no competing interest.

